# Elucidating Heterogeneous Iron Biomineralization Patterns in a Denitrifying As(III)-Oxidizing Bacterium: Implications for Arsenic Immobilization

**DOI:** 10.1101/2022.01.28.478188

**Authors:** Rebeca Lopez-Adams, Simon M. Fairclough, Ian C. Lyon, Sarah J. Haigh, Jun Zhang, Fang-Jie Zhao, Katie L. Moore, Jonathan R. Lloyd

**Affiliations:** Department of Earth and Environmental Sciences, University of Manchester, Manchester, UK; Department of Materials, University of Manchester, Manchester, UK; Department of Materials Science and Metallurgy, University of Cambridge, Cambridge, UK; College of Resources and Environmental Sciences, Nanjing Agricultural University, Nanjing, China; Photon Science Institute, University of Manchester, Manchester, UK

## Abstract

Anaerobic nitrate-dependent iron(II) oxidation is a process common to many bacterial species, which promotes the formation of Fe(III) minerals that can influence the fate of soil and groundwater pollutants, such as arsenic. Herein, we investigated simultaneous nitrate-dependent Fe(II) and As(III) oxidation by *Acidovorax* sp. strain ST3 with the aim of studying the Fe biominerals formed, their As immobilization capabilities and the metabolic effect on cells. X-ray powder diffraction (XRD) and scanning transmission electron microscopy (STEM) nanodiffraction were applied for biomineral characterization in bulk and at the nanoscale, respectively. NanoSIMS (nanoscale secondary ion mass spectrometry) was used to map the intra and extracellular As and Fe distribution at the single-cell level and to trace metabolically active cells, by incorporation of a ^13^C-labeled substrate (acetate). Metabolic heterogeneity among bacterial cells was detected, with periplasmic Fe mineral encrustation deleterious to cell metabolism. Interestingly, Fe and As were not co-localized in all cells, indicating delocalized sites of As(III) and Fe(II) oxidation. The Fe(III) minerals lepidocrocite and goethite were identified in XRD, although only lepidocrocite was identified via STEM nanodiffraction. Extracellular amorphous nanoparticles were formed earlier and retained more As(III/V) than crystalline “flakes” of lepidocrocite, indicating that longer incubation periods promote the formation of more crystalline minerals with lower As retention capabilities. Thus, the addition of nitrate promotes Fe(II) oxidation and formation of Fe(III) biominerals by ST3 cells which retain As(III/V), and although this process was metabolically detrimental to some cells, it warrants further examination as a viable mechanism for As removal in anoxic environments by biostimulation with nitrate.

## 1. Introduction

Arsenic is a toxic element whose solubility and toxicity are governed by chemical speciation. This element is mobilized in the environment through anthropogenic activities, such as mining and agriculture, or through natural processes such as volcanic activity, weathering of As-bearing minerals and through microbial processes^1^. Bacteria capable of metabolising As are found in diverse ecosystems and are acknowledged as playing a key role in mobilizing As in the environment, particularly through the reductive dissolution of As(V)/Fe(III) mineral assemblages^2-5^. Arsenic has a high affinity for sulfur and iron minerals in the subsurface, therefore, the formation of new mineral phases containing these elements could stimulate the immobilization of As in the environment^6^. Denitrification is a widespread microbial process, particularly in bacteria, where nitrate is enzymatically reduced to N_2_ in the absence of oxygen^7^. During denitrification, nitrate is used as a terminal electron acceptor coupled to the oxidation of suitable electron donors, such as arsenite [As(III)]^8-10^, which some bacteria can oxidize to arsenate [As(V)], the generally assumed less toxic and less mobile As oxyanion^11^. In addition, some heterotrophic denitrifying bacteria are able to catalyze Fe(II) oxidation, in a process known as nitrate-dependent iron(II) oxidation (NDFO), which is thought to be a result of both biotic and indirect biotic effects^12^, although, the mechanism of Fe(II) oxidation in these heterotrophs is not fully resolved, and no enzymatic Fe(II) oxidation has been reported to date^13, 14^. Moreover, nitrite and nitric oxide can abiotically oxidize Fe(II). These reactive nitrogen species are also intermediary compounds produced and accumulated during denitrification^15^, and have been shown to be responsible for most of the Fe(II) oxidation observed^16, 17^. Organics and particularly extracellular polymeric substances (EPS), produced by many NDFO bacteria, have also been suggested as important role players in Fe(II) oxidation. EPS contain polysaccharides that are known to complex Fe(II), therefore, EPS could also be a site for mineral precipitation^18^.

Fe(II)-oxidizing bacteria have been reported to produce the intermediate oxidation state Fe minerals green rust^18, 19^ and magnetite^19-22^, and Fe(III)-oxyhydroxides such as ferrihydrite^23^, goethite^15, 24, 25^ and lepidocrocite^24-27^. Additionally, NDFO bacteria can precipitate these Fe minerals at different cell sites, including the periplasm and the cell surface^28-34^. The periplasm is a crucial cellular compartment for energy generation and the transit of nutrients and waste, therefore, its encrustation with minerals is thought to be lethal to the cells^35^. Furthermore, it has been proposed that Fe(II) oxidation in anaerobic denitrifying conditions is a detoxification mechanism rather than an energy conserving reaction^35-37^, where the Fe(II) concentration in these incubations is typically higher than in environmental conditions and could be the reason for this cell mineralization^38^.

Many electron microscopy and spectroscopic techniques have been used to study the NDFO process and biomineralization patterns in bacteria^38-42^, more extensively in *Acidovorax* sp. strain BoFeN1^15, 24, 29, 32, 43^. These studies report the distinctive Fe mineral phases produced depending on the bacterial strain and the physical, geochemical and growth medium conditions. For instance, a high pH medium with added carbonate has been observed to favour the formation of goethite^24^. The presence of As(III/V) in the growth medium has also been tested, and arsenic may alter the crystallinity of the biominerals formed under certain conditions^23^.

The facultative bacterium *Acidovorax* sp. strain ST3, with a capacity to oxidize As(III) upon the addition of nitrate, was isolated from a paddy soil polluted with arsenic, where it was hypothesised that As(III) oxidation was coupled to nitrate reduction^10^. In this initial work, the periplasmic AioAB arsenite oxidase was identified and this bacterium was also described as a NDFO organism^10^. In the present work, the simultaneous oxidation of As(III) and Fe(II) by *Acidovorax* sp. strain ST3 under denitrifying conditions was investigated. The aim was to identify the composition and precipitation site of the Fe biogenic minerals produced by strain ST3 and assess their capability to immobilize As. Isotopic labelling and NanoSIMS single-cell depth profiling were used to assess the effect of As and Fe in cells, to locate the site(s) of mineral precipitation and to infer where Fe(II) oxidation occurs. Additionally, transmission electron microscopy (TEM), STEM, energy dispersive X-ray spectroscopy (EDS), electron energy loss spectroscopy (EELS) and selected area electron diffraction (SED) were performed to characterize the biominerals chemically and structurally at the nanoscale. It was hypothesized that ST3 cells would catalyze As(III) oxidation to As(V), and the newly formed Fe(II/III) mineral phases would be thermodynamically stable hosts for As(V). Understanding the nano-scale mineralogy of iron and arsenic biooxidation is therefore important to help understand the long-term immobilization of As in natural environments by this process. The greater knowledge of the fundamentals of biooxidation will support the application of As(III)-oxidizing bacteria to remediate As(III)-polluted environments.

## 2. Materials and methods

### 2.1 Culture conditions

*Acidovorax* sp. strain ST3 was first streaked onto aerobic lysogeny broth (LB) agar plates and incubated for 48 h at 28°C in the dark. Single isolated colonies were transferred to Erlenmeyer flasks containing 100 mL liquid LB broth and incubated in a shaker/incubator at 28°C/150 rpm for 24 h. This liquid culture (5 mL) was used to inoculate serum bottles containing 100 mL of anoxic low-phosphate (LP) medium^44^. These bottles were incubated for 48 h/28°C in the dark (until late logarithmic phase), and this culture was used as inoculum for metal oxidation experiments. The cultures were transferred to 50 mL conical tubes and centrifuged (2509 g/30 min) for harvesting. The pellet was washed twice with 20 mL of 10 mM anaerobic piperazine-N, N′-bis(2-ethanesulfonic acid) (PIPES) buffer (pH=7.0) and centrifuged (2509 g/30 min). The last pellet was resuspended in PIPES buffer (3-5 mL). These processes were performed under a N_2_/CO_2_ (90:10 v/v) atmosphere. All the serum bottles used to prepare anoxic medium, buffers and the experiments were degassed with N_2_ (>30 min) prior to the transfer of media or buffer solutions.

### 2.2 Anaerobic Fe(II) and As(III) oxidation experiments

Washed cells of strain ST3 were incubated in serum bottles containing 20 mL of anoxic LP medium and adding 10 mM FeCl_2_, 10 mM ^13^C-labelled sodium acetate (^13^CH_3_CO_2_Na) to label metabolically active cells, and 0.5 mM As(III) (NaAsO_2_, Sigma-Aldrich^®^), which was added after filtration of the medium to remove a green-whitish Fe(II) precipitate, as described elsewhere^15^. After filtration, the final concentration of Fe(II) was 3.5-4.0 mM.

Two growth conditions were tested, planktonic and biofilm, depending on the analysis techniques (planktonic for XRD and STEM, and biofilm for scanning electron microscopy and NanoSIMS). For planktonic experiments, the bottles with LP medium were inoculated with ST3 cells (final OD_600_≈0.15). For biofilm experiments, two boron-doped silicon (Si) wafers (7.2 × 7.2 × 0.5 mm) per experimental bottle were introduced vertically in a plastic holder, the plastic holder was fixed to the bottom of the bottle with silicon grease, after which the bottles were sealed and degassed with N_2_ prior to the addition of medium and cells by injection. The Si wafers supported the formation of a biofilm and mineral precipitates (see Figure S1 C). The experiments were carried out in triplicate and three additional bottles of each growth condition were left un-inoculated (“no cell” controls). All the experiments and controls were incubated at 28°C in the dark for 14 days (biofilm) and up to 21 days (planktonic).

### 2.3 Chemical analysis

The aqueous phase was sampled (100 µL) every 2-3 days and filtered immediately (0.22 µm pore size filters, Jet Biofil^®^). These filtered samples were analyzed for Fe(II), As species, total Fe (Fe_total_), total As (As_total_), nitrate, nitrite and acetate. The ferrozine method was used to measure Fe(II)^45^. Arsenic species samples were diluted in distilled H_2_O and quantified by inductively coupled plasma-mass spectrometry (ICP-MS, 7500CX Agilent Technologies, USA)^46^. The anions nitrate, nitrite and acetate were quantified by ion chromatography using a Dionex ICS 5000 equipped with an AS11HC 0.4 mm capillary ion exchange column. Fe_total_ and As_total_ samples were extracted in 2 % HNO_3_ and analyzed in ICP atomic emission spectroscopy (ICP-AES) using a Perkin–Elmer Optima 5300 DV instrument.

### 2.4 SEM and NanoSIMS imaging

A FEI Quanta 650 field emission gun scanning electron microscope (FEG SEM) operating at 15 kV, using the secondary electron (SE) detector under high vacuum, was used to find suitable areas prior to NanoSIMS analysis and to assess the effect of the sample preparation. Samples were subsequently analyzed in a NanoSIMS 50L ion microprobe (CAMECA, France) using a Cs^+^ primary ion beam at 16 keV. A beam with a current of 0.131-1.435 pA with spatial resolution from 150-300 nm (D1 aperture= 4-2) was scanned over the surface of the sample and the secondary ions were collected and analyzed using a double focusing mass spectrometer. The following negative secondary ions were detected simultaneously in multicollection mode: ^12^C, ^13^C, ^12^C^14^N, ^28^Si, ^56^Fe^12^C, ^56^Fe^16^O and ^75^As. Additionally, an ion-induced SE image was obtained. Prior to the start of collection, the regions of analysis (ROIs) were implanted with Cs^+^ ions to a dose of 1×10^17^ ions cm^-2^ to remove the Pt coating (10 nm) and reach steady state^47^. Images were collected at a dwell time between 2000-5000 µs px^-1^. The CAMECA mass resolving power (MRP) was >7000 using ES=3 and AS=2. CAMECA high mass resolution spectra were acquired at masses 13, 72 and 75 using iron metal and gallium arsenide as reference materials to avoid peak overlaps due to molecular interferences, primarily ^12^C^1^H at mass 13 (^13^C), ^28^Si_216_O at mass 72 (^56^Fe^16^O) and ^56^Fe^19^F at mass 75 (^75^As)^48^. The image pixel sizes were 128 × 128 or 256 × 256. Further information on sample preparation and analysis can be found in the Electronic Supplementary Information (ESI) (Text S1.1 and S1.4).

#### 2.4.1 NanoSIMS depth profiles using Cs^+^ or O^-^ primary ions

After collecting nanoSIMS ions images with large fields of view (30-60 µm to image groups of cells and minerals), single cells were selected to collect image depth profiles with smaller square raster sizes (3-7 µm width). Both Cs^+^ (for producing negative secondary ions) and O^-^ (for producing positive secondary ions) were used as primary ions for depth profiling and the selected areas were not implanted with Cs^+^ or O^-^ ions prior to starting the analysis, to preserve and probe the intact biomineral coating on the cells. The spatial resolution was improved by using D1=4 or 5 (beam size=120-100 nm). For details of the depth profile set-up and NanoSIMS data analysis see the ESI Text S1.5.

### 2.5 Biomineral characterization

#### 2.5.1 Analytical TEM, STEM, EDS and EELS

Planktonic samples were analyzed by transmission electron microscopy (TEM) in bright field (BF) imaging mode using a JEOL F200 with an ASTAR/Digistar-Topspin nanodiffraction setup and also by high-angle annular dark-field (HAADF) STEM (for higher resolution imaging) with a FEI Titan G2 ChemiSTEM at 200 kV using a 180 pA beam current. The Titan instrument was also used to collect dual EELS and EDS data simultaneously using the SuperX EDS detector system and the Gatan GIF Quantum ER spectrometer via Gatan’s Digital Micrograph software. The EDS and EELS data were then processed using Hyperspy. EDS quantification was achieved using background correction and a standardless Cliff Lorimer analysis, whilst EELS quantification of the Fe^3+^ to Fe^2+^ ratio was calculated using the relative intensities of the Fe L2, L3 peaks as described elsewhere^49^. This processing allowed the collection of high-resolution elemental maps of the biominerals. Arsenic does not have well defined core loss peaks, therefore local differences in As oxidations states cannot be distinguished. The apparent nanoparticles (NPs) size measured from BF TEM images was estimated manually with ImageJ. For further details on sample preparation and anoxic sample loading see the ESI Text S1.2 and S1.3.

#### 2.5.2 Powder X-ray diffraction (XRD)

Bulk biominerals were identified in an XRD Bruker^©^ D8 Advance X-ray diffractometer, where the slides were placed in an airtight specimen holder (Bruker©) to maintain anoxic conditions. The samples were analyzed with a 40 kV beam, over the 5-70° 2θ range, with a 0.02° 2θ step size and 0.5 s counting time. Spectra were processed with XRD DIFFRAC.EVA software (Bruker^©^).

## 3. Results and discussion

### 3.1 Aqueous geochemistry

Fe(II) oxidation was noticeable as the formation of rusty precipitates from day 1 in the experimental bottles containing cells (Figure S1 A). Both planktonic and biofilm culturing conditions favoured Fe(II) oxidation to a similar rate and extent, although biofilm conditions slightly enhanced this oxidation. At day 7 the biofilm samples had oxidized 67% of the Fe(II) to Fe(III), while the planktonic samples oxidized 62% of Fe(II) (Figure S2 A), as indicated by the ferrozine analysis. By day 14 the biofilm samples oxidized 72% of Fe(II), whereas the planktonic samples oxidized 65% of Fe(II) (Figure S2 A). Only planktonic samples were monitored until day 21 when 81% of Fe(II) was oxidized (Figure S1 A). There was a removal of 15 % of the Fe(II) by day 7 in the “no cells” control, and over the course of 21 days this abiotic Fe(II) removal increased to 31 % (Figure S2 A). These control samples presented a reddish layer at the surface of the medium (Figure S1 B), suggesting that some oxygen might have been introduced, perhaps during sampling, which caused abiotic Fe(II) oxidation, but no visible precipitates were formed. Additionally, further abiotic reaction of Fe(II) with carbonate and phosphate from the growth medium was possible^25^.

The aqueous As speciation analysis showed a removal of 93 % As(III) in biofilm samples and a 83 % As(III) removal in planktonic samples at day 7 (Figure S2 B). As(V) was detected in all experiments from day 1 and dropped in concentration over the duration of the experiment (from 110 to 10 µM) (Figure S2 C). There was also 66 % As(III) removal in the “no cells” controls (Figure S2 B), likely due to abiotic Fe(II) oxidation, as described previously. The As_total_ measurements by ICP-AES were below level of detection (10 ppb) in both biofilm and planktonic experiments after 7 days, whereas the “no cells” controls showed 75% removal of As_total_ (Figure S2 D). As(V) concentrations did not increase with incubation time in the aqueous phase, in contrast to a previous report of this strain being able to couple anaerobic As(III) oxidation to nitrate reduction^10^. As(V) was most likely sorbed to the solid biominerals, which were removed from the aqueous phase by filtration prior to the analysis of As species. Moreover, spontaneous oxidation of As(III) by exposure to oxygen would not explain the soluble As(V) quantified during all time points, including at the start of the experiment, as this process is much slower^50^.

The consumption of acetate and nitrate was followed until day 7 in all experiments, where 0.61 mM of acetate were consumed in the biofilm samples and 0.26 mM was consumed in planktonic samples, implying a 5.5% and 2.5% consumption of acetate in the biofilm and planktonic samples, respectively (Figure S3 A). Acetate was added in excess in this work, however, Fe(II)-oxidizers have been found to only require organic co-substrates at low concentrations, between 0.5 to 1.0 mM^38^. 1.99 mM of nitrate was consumed in biofilm and 0.99 mM in planktonic conditions, indicating that 17 % and 9 % of nitratewas consumed in biofilm and planktonic growth samples, respectively (Figure S3 B). Nitrite was not detectable in the supernatants over the duration of the experiment, which contrasts with other NDFO bacteria^10, 15, 27, 40^. However, considering the low nitrate consumption, it is possible that nitrite was produced at much lower concentrations and immediately consumed by reaction with Fe(II)^51^. The “no cells” control showed no changes in the acetate and nitrate concentrations, and nitrite was not detected either, as expected.

Overall, biofilm conditions were more favourable for As(III) removal, Fe(II) oxidation and denitrification. Bacteria are commonly found growing in biofilms in natural systems. Biofilms contain metal-reactive ligands, for instance in the EPS fraction, which could have complexed Fe(II) and promoted its precipitation^52^ and subsequent sequestration of As(III/V). Additionally, the Si wafer surface in biofilm samples might have played a role as a nucleation site for biominerals, precipitating Fe and As, although not necessarily as the oxidized species.

### 3.2 Imaging of As and Fe using SEM and NanoSIMS

SEM images of biofilm samples showed abundant wafer surface colonization in all time points analyzed, and many cells were coated with minerals (‘mineralized cells’) from day 4 of incubation (not shown). The chemical fixation-dehydration treatment proved better than the air-dried method to preserve the shape and structure of cells, and thus more suitable for NanoSIMS analysis (see ESI Text S2 & Figure S4). Samples from days 7 and 14 were selected for further NanoSIMS analysis (Figures S5 & 1, respectively). At day 7 many cells showed high Fe counts (as ^56^Fe^16^O), indicating that an Fe mineral was coating the surface of cells (Figure S5 A-C). Extracellular biominerals (attached or detached from the cells) presented higher ^56^Fe^16^O counts on their surfaces than the cells. ^75^As was co-located with ^56^Fe^16^O on the surfaces of both cells and extracellular biominerals (seen in orange in Figure S5 A-C), although ^75^As was not detected in all mineralized cells. The cell poles showed higher ^56^Fe^16^O counts than on the rest of the cell, suggesting preferential mineral encrustation in this region (Figure S5 C). This was previously observed in strain BoFeN1 ^32^ and was later confirmed herein by NanoSIMS depth-profiles (Figure S6). By day 14, some cells showed ^56^Fe^16^O counts on their surface (white arrows Figure 1A-C) with comparable intensity as the ^56^Fe^16^O counts on the extracellular biominerals (pink arrows Figure 1A-C), suggesting heavier Fe mineralization of the cells after this prolonged incubation period. The surface of mineralized cells at both time points presented a higher intensity signal for ^56^Fe^16^O than for ^56^Fe^12^C, despite the abundance of C in cells, where even the ^12^C signal was low. This could indicate that only the extracellular mineral coating covering the cells was probed by the Cs^+^ beam and the primary ions did not reach the cell surface. Alternatively, ^56^Fe^16^O could be a molecular ion with higher probability of formation under these analytical conditions. The data presented are consistent with the presence of an Fe (oxyhydr)oxide mineral coating on the cells.

**Figure 1.**
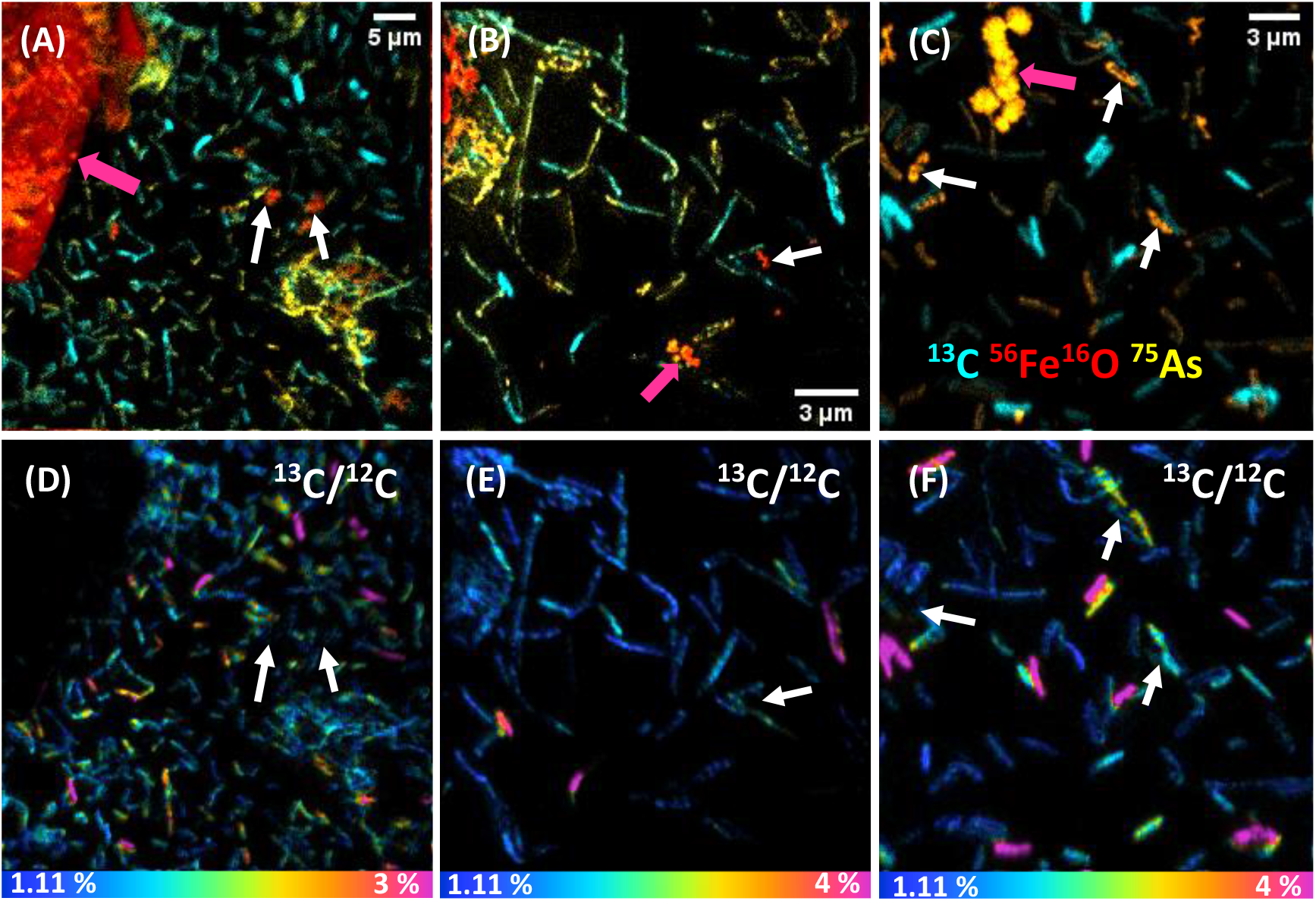
NanoSIMS images of biofilm ST3 cells after 14 days of incubation and preserved by chemical fixation and dehydration. The top row (A), (B) and (C) are overlay images displaying ^13^C (cyan), ^56^Fe^16^O (red) and ^75^As (yellow). The bottom row (D), (E) and (F) are the ^13^C/^12^C HSI images of (A), (B) and (C), respectively. Note the heterogeneous ^13^C accumulation by cells in (D)-(F). White arrows in (A)-(C) are heavily mineralized cells that do not show any ^13^C signal (corresponding white arrows in (D)-(F)). Pink arrows in (A)-(C) indicate Fe-biominerals of different size and morphology. The horizontal color bars in (D)-(F) are the ^13^C % accumulation, where blue is natural abundance (1.11 %).

### 3.3 Metabolic heterogeneity revealed at the nanoscale and correlation with cell mineralization

By day 7 the cells showed negligible ^13^C assimilation (Figure S5 D-F), in agreement with the low levels of aqueous acetate consumption. However, some regions in these cells showed up to 2% ^13^C/^12^C assimilation, and interestingly, the parts of the cells that were not ^13^C-enriched showed ^56^Fe^16^O counts (Figure S5 C & F). Moreover, many cells were fully coated with mineral, and showed no ^12^C or ^13^C signal (Figure S5 B & E). In contrast, by day 14 cells showed heterogeneous but higher ^13^C labelling, assimilating up to 4% of ^13^C/^12^C (Figure 1 D-F), with a mean content of 2.6 ± 1.3%. Thus, cells assimilated on average 2.3 times more ^13^C than natural abundance (Figure 2), signifying a culture metabolic recovery, distinct from the first 7 days of incubation. However, many cells were not metabolically active and these had a ^13^C/^12^C ratio closer to the natural abundance. Most cells that were mineralized were not metabolically active, but presented an intense ^75^As signal, suggesting As sorption to the Fe mineral (Figure 1A-C). Interestingly, most active cells did not show ^56^Fe^16^O on the surface but ^75^As was associated with some of these cells, suggesting that it was surface-bound^53^ or taken up intracellularly. Only a few cells showed simultaneous Fe mineralization, ^75^As on the surface and ^13^C assimilation.

**Figure 2.**
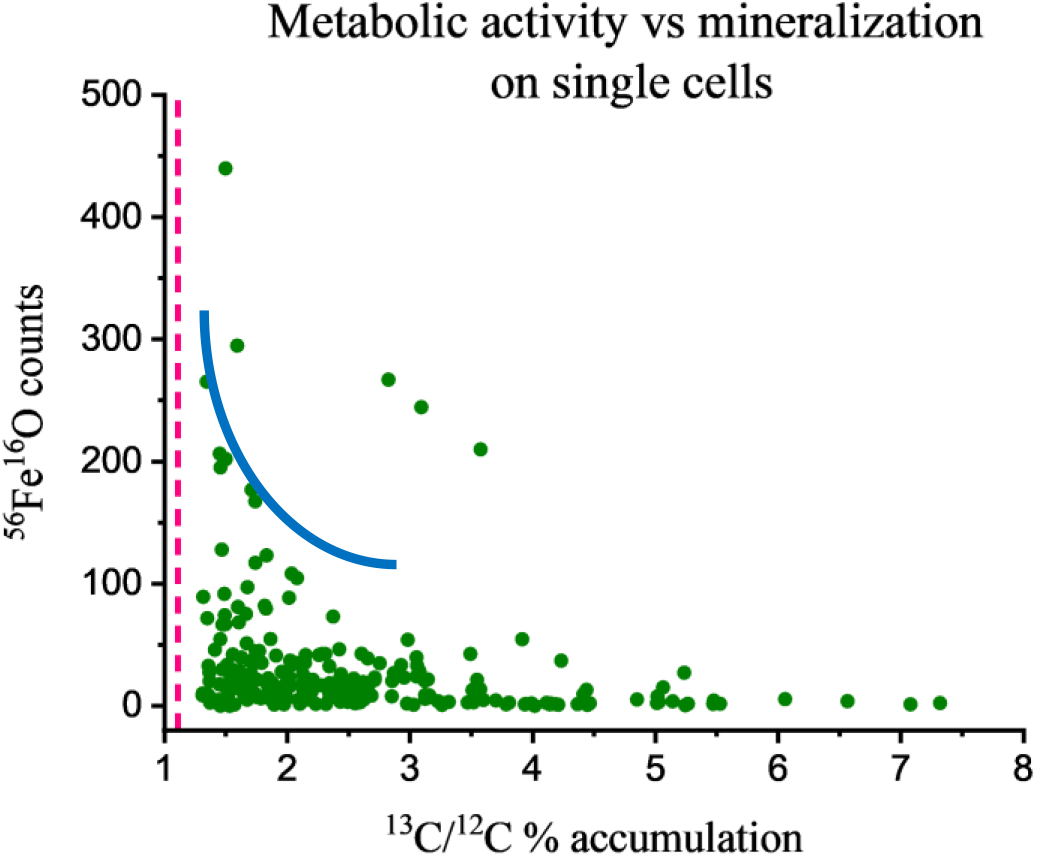
NanoSIMS ^56^Fe^16^O counts (mineralization) versus ^13^C/^12^C % (metabolic activity) on single cells in biofilm conditions at day 14 of incubation. Cells only accumulate high levels of ^13^C at low levels of ^56^Fe^16^O (low mineralization), although most cells are in a region in the plot (blue curved line) where low mineralization is compatible with moderate ^13^C accumulation. The pink vertical dashed line is the natural ^13^C/^12^C abundance, 1.11 %. N=230.

The correlation between metabolic activity and the level of cell mineralization can be estimated from the measured ^56^Fe^16^O counts at the cell surface (normalized to ion dose) and the ^13^C/^12^C ratio (Figure 2). Broadly this shows a negative correlation between metabolic activity and Fe mineralization at the single cell level. This indicates that metabolically active cells can deter cell mineralization, or most likely, high cell mineralization inhibits the metabolic activity of ST3 cells, an effect previously observed in strain BoFeN1^35^. From these results, it can be inferred that cell encrustation was metabolically disadvantageous. This Fe mineralization probably blocked the flow of substrates inside and outside of the cell and disturbed the enzymatic oxidation of As(III) and reduction of nitrate. ST3 cells can tolerate high concentrations of arsenite (5 mM)^10^, however, it is unclear if cell mineralization was a protection mechanism against Fe(II)^38^, added at five times higher the concentration than in the soil porewater where ST3 was isolated, or if it was a harmful effect of Fe(II) in combination with high concentrations of As(III) (>400 µM). This co-toxicity was previously reported for Fe(II), which increased in toxicity in the presence of Cu(II)^54^.

The heterogeneous pattern of Fe mineralization and ^13^C accumulation indicates an interesting phenotypic diversity in genetically identical cells cultured under the same conditions. Metabolic heterogeneity is an ongoing research field, where not only the genotype and environment regulate these variations but also stochastic gene expression^55, 56^. In microbial populations, ecological factors such as surfaces or cell-to-cell communication, the cell cycle and cell ageing can affect gene expression, leading to metabolic variations. The cell cycle and cell ageing might explain the different biomineralization patterns observed in this work, which is expected under batch culture conditions^35^. For instance, cells could be at different growth stages, where some cells may be reducing nitrate and producing nitrite, and those producing more nitrite could quickly form Fe(III) precipitates, blocking the periplasm and inhibiting further uptake of ^13^C-labelled organics. Additionally, the marked mineralization at the (older) cell poles could be explained by protein aggregation in this part of the cells, as proposed earlier^35^.

### 3.4 Spatial distribution of As and Fe at the single-cell level

NanoSIMS single-cell depth profiles were obtained to evaluate the As and Fe distribution in whole cells at high resolution, where 3D reconstructions facilitated the visualisation of the relative distribution of As and Fe (Figures 3 & S6). Many ST3 cells appeared to be covered in Fe minerals during the initial NanoSIMS imaging, and this mineralization indicated that ^75^As co-located with ^56^Fe^16^O in most cells (Figures 1 & S5). However, a more detailed analysis by 3D remodelling revealed asymmetric Fe encrustation in cells, where surprisingly, ^75^As and ^56^Fe^16^O were not spatially co-located in all cells (Figure 3A & B). Regardless of the level of cell encrustation, ^75^As was imaged on all cell surfaces (see arrow 1 in Figure S5 A), and cells that had higher metabolic activity showed ^75^As but no ^56^Fe^16^O at the surface (see bottom cell in Figure 3A-E and arrow #1 in Figure S6 A). ^75^As was not detected intracellularly and was presumably located at the periplasm (Figures 3A-B & S6 A-B). This could be indicative of As(III) binding proteins in this region^53^, and the ^75^As observed to be co-locating with encrusted Fe could be mixed valence As, given that Fe minerals can sorb both As(III) and As(V)^57^. The 3D reconstructions showed this asymmetrical As and Fe distribution at the cell surface and periplasm, in contrast to the classical NanoSIMS stack images (panels C, D, H & I in Figures 3 & S6), which only show the total sum of each secondary ion in 2D, masking this asymmetry. Moreover, the cell poles showed a more pronounced encrustation observed under Cs^+^ ion bombardment (Figure S6 B), which was not observable in O^-^ ion bombardment, where the whole cells appeared encrusted (Figures 3F-G & S6 F-G). However, this technique artefact is acknowledged, because ^56^Fe ionizes more easily under O^-^ than under Cs^+^ bombardment, producing a higher intensity signal. Regardless of the secondary ion collection mode, the ^56^Fe^+^/^56^Fe^16^O^-^ signals were more intense than ^75^As in depth profiles, suggesting a higher abundance of Fe localized within or at the surface of the cells.

**Figure 3.**
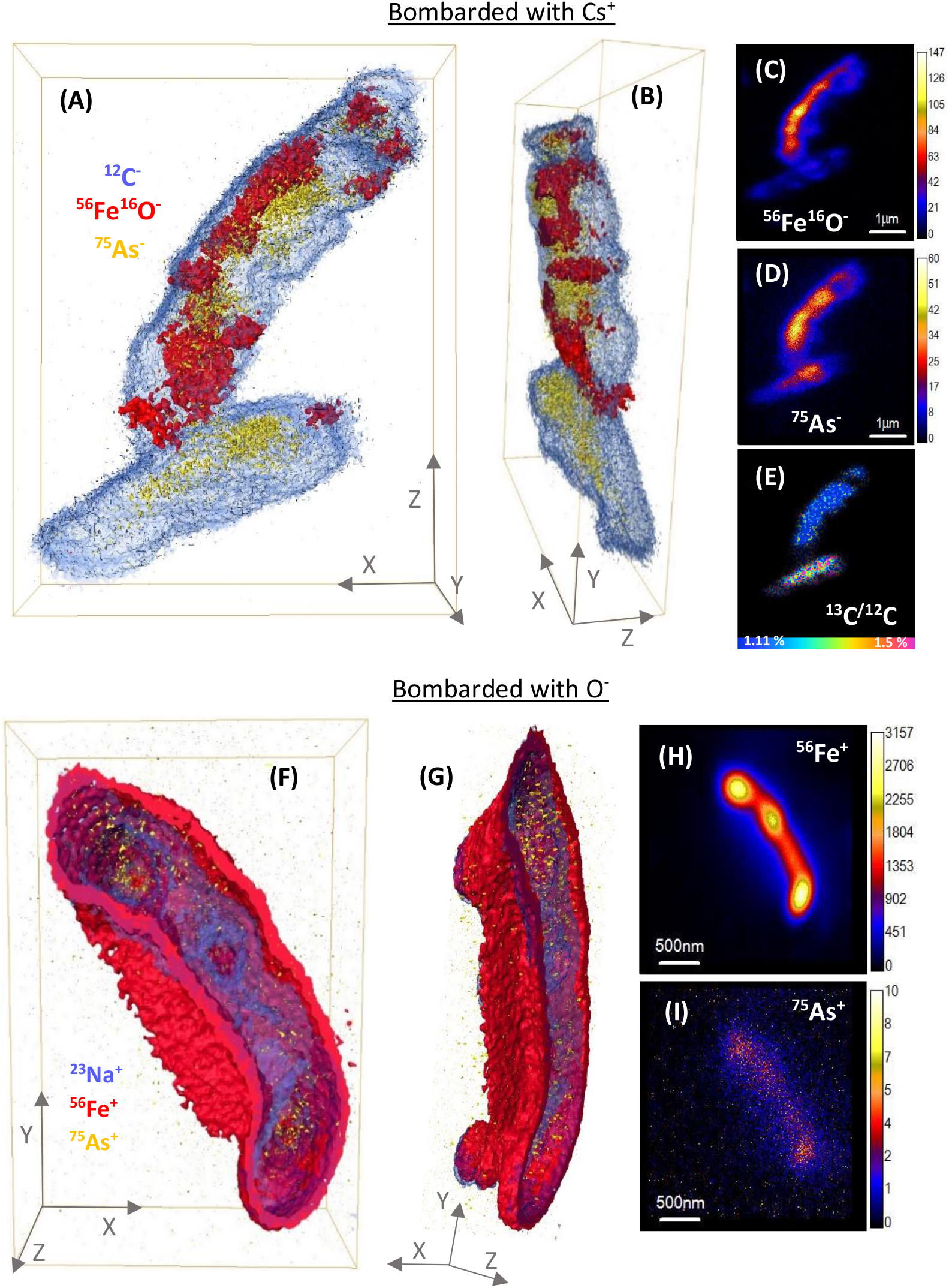
3D reconstructions of NanoSIMS depth profiles of strain ST3. (A), (B), (F) and (G) are 3D models, where (A) are cells sputtered with Cs^+^ and (B) is the side view of the same cells; (F) is a cell sputtered with O^-^ and (G) is the side view of the same cell. In these 3D reconstructions (A, B, F & G) Fe is shown in red (either ^56^Fe^16^O^-^ or ^56^Fe^+^), ^75^As in yellow and ^12^C^-^ (A & B) or ^23^Na^+^ (F & G) in blue. (C) and (D) are stack images of ^56^Fe^16^O and ^75^As, respectively, of the same cells in (A); (E) is the ^13^C/^12^C ratio of the same area, sum of 74 planes. (H) and (I) are stack images of ^56^Fe and ^75^As, respectively, of the cell in panel (F), sum of 71 planes. The vertical color bars in panels (C), (D), (H) & (I) are the secondary ion intensities.

Prior to this work, mineralized BoFeN1 cells had been analyzed in 3D via nanoscale techniques, and different Fe biomineralization patterns (e.g. encrusted periplasm or spike-shaped iron minerals) were reported^29^. However, the study did not distinguish metabolically active and inactive cells. To our knowledge, the present work is the first to use NanoSIMS imaging to generate 3D models of microbial biomineralization patterns in metabolically active or inactive nitrate-dependent Fe(II)-oxidizing cells, in simultaneous incubations with As(III).

### 3.5 Mineral identification using powder X-ray diffraction (XRD)

The precipitates obtained by planktonic growth were analyzed by powder XRD. The spectrum from the sample at day 7 showed no significant peaks, indicating the presence of a poorly crystalline Fe phase, although by day 10 the more crystalline phases lepidocrocite (γ-Fe^3+^OOH) and goethite (α-Fe^3+^OOH) were identified, which showed broad but higher intensity peaks in the air-exposed analysis than during anaerobic analysis (Figure S7). These Fe(III)-oxyhydroxides have been reported as a product of NDFO bacteria grown in medium with carbonate^24^.

### 3.6 Crystallinity and biomineral identification at the nanoscale with TEM

The planktonic precipitates were imaged in TEM after 1, 3 and 7 days of incubation, to observe the early formation of cell-associated/detached biominerals. The cells showed mineral encrustation from day 1, where extracellular biominerals were present. The spherical biomineral nanoparticles (NPs) showed no crystallinity and were amorphous at the cell surface or detached from the cells (Figure 4A & B). The cells showed heavier mineralization at the poles than in other regions of the cells (Figure 4A-B). By day 3, the coating had thickened, especially at the polls, with the NPs having similar morphology to that seen for day 1 (Figure 4C). By day 7, similar mineralization and morphologies were observed, but additionally a new phase was noted on the surface of the cells. This new biomineral appeared to have a more crystalline structure, resembling “flakes” (Figure 4E-F). The NP size increased from a mean of 48 ± 25 nm at day 1 to 104 ± 43 nm by day 7 (errors represent standard deviation). Moreover, regardless of the time point, the NPs formed aggregates detached from the cells (Figure 4D) or aggregated as “filaments” associated to the cell surface (Figure 4A, C & E), suggesting a possible effect of EPS that acted as mineral nucleation sites^58, 59^. By day 7 the detached NPs were twice as big as those on the cell surface (Figure 4E).

**Figure 4.**
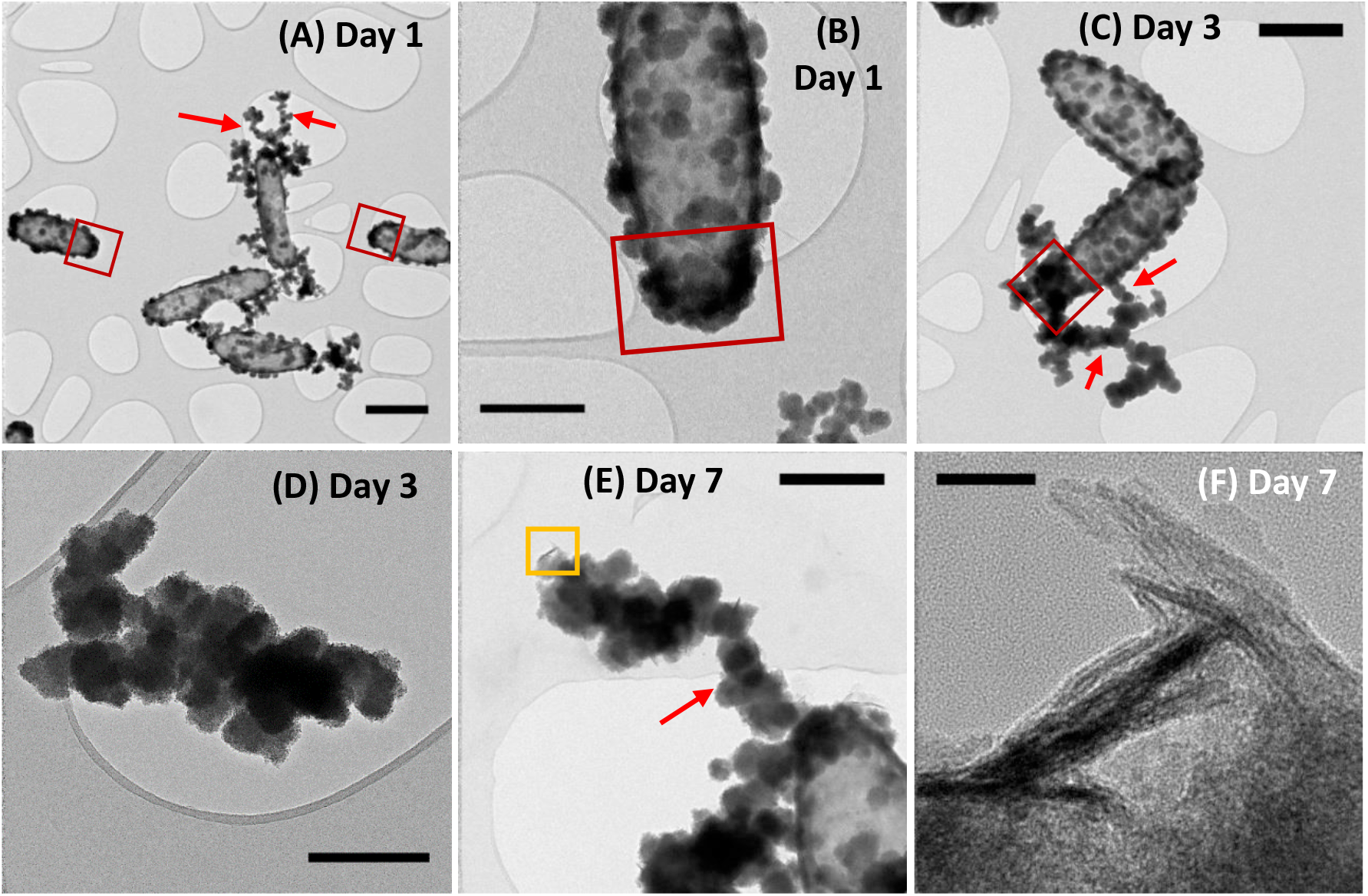
Bright field TEM micrographs of planktonic ST3 samples at days 1 (A and B), 3 (C and D) and 7 (E and F). Mineralized cells can be observed from day 1, showing a thicker mineralization by day 3 and the appearance of a more crystalline structure by day 7. The red squares in (A)-(C) highlight the cell poles with heavier mineralization than the rest of the cell. The yellow square in (E) is a selected region with “flake-like” biominerals at higher magnification in panel (F). Red arrows in A, C & E are indicating the nanoparticles aggregated as filaments associated with the cell surface. Scale bars are 500 nm in (A), (B), (C) and (E), 300 nm in (D) and 20 nm in (F).

### 3.7 High-resolution elemental mapping of biominerals using STEM-EDS

Both biomineral morphologies identified at day 7 (crystalline “flakes” and amorphous spherical NPs) were further analyzed using STEM-EDS and EELS (Figure 5), with high-resolution data collected from the outer (thinner) regions of these biominerals. All the biominerals were mainly composed of iron, oxygen and arsenic, irrespective of their morphology, and their atomic abundance was quantified (Table S1).

**Figure 5.**
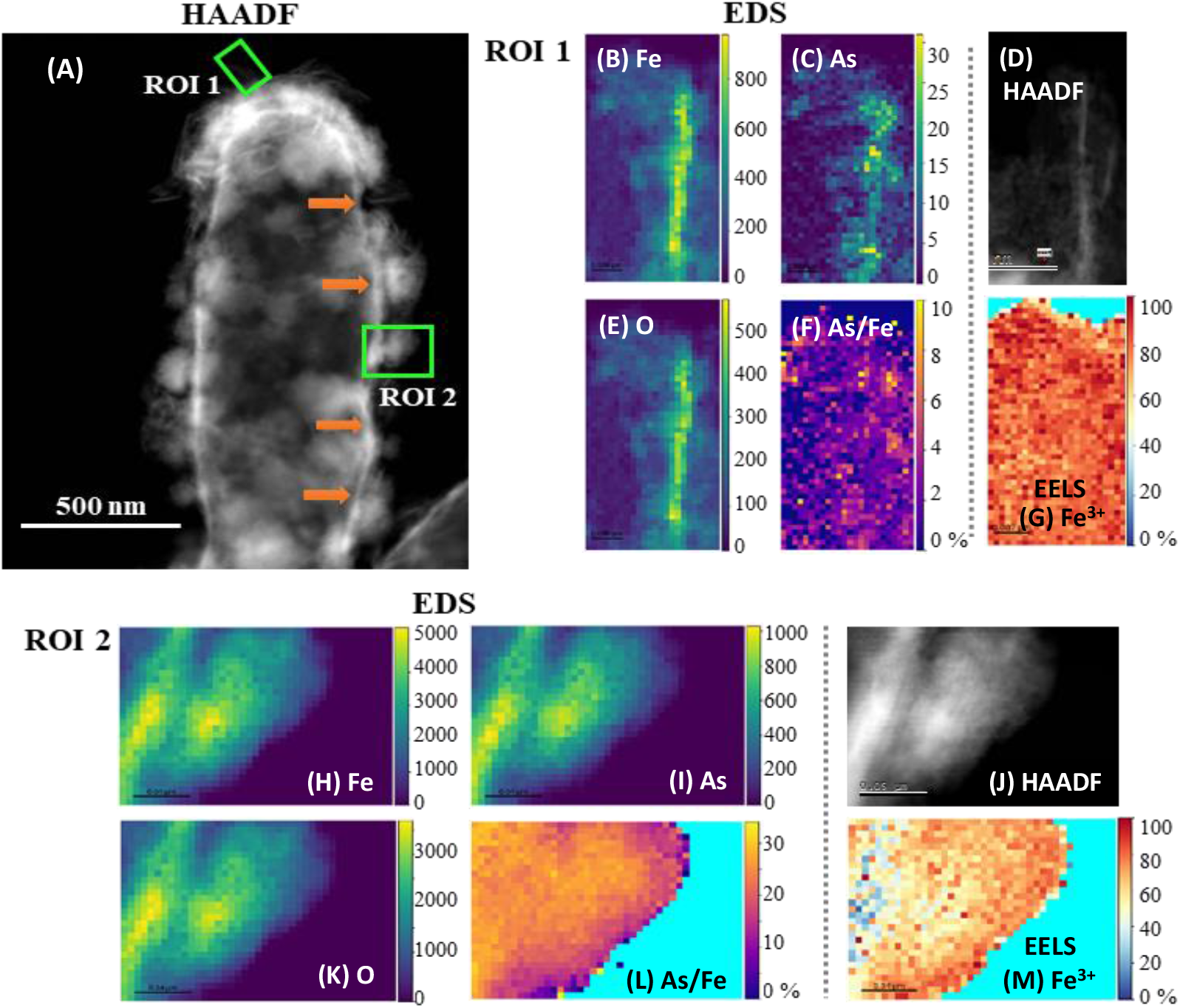
High-angle annular dark field (HAADF) STEM micrographs with complementary EDS/EELS analysis of two regions in a mineralized ST3 cell. (A) is the HAADF-STEM micrograph of the whole cell indicating the selected ROIs 1 and 2 (green squares). EDS maps of Fe, As and O in ROI 1 are shown in (B), (C) & (E), respectively, and in (H), (I) & (K) for ROI 2, where the vertical colour scale bars show the atomic count intensities. (F) & (L) are atomic As/Fe ratio (%) maps of ROIs 1 and 2, respectively. (D) & (J) are HAADF images, and (G) & (M) are EELS Fe^3+^ maps of regions 1 and 2, respectively. Fe, As and O co-located strongly in the amorphous biomineral in ROI 2, whereas only Fe and O co-localized strongly in the “flake” in ROI 1. The atomic As/Fe ratios (atomic %) show that the mean As/Fe was 2.4 % in ROI 1 and 23.4 % in ROI 2, suggesting that the amorphous biomineral in ROI 2 retained more As than the crystalline “flake” in ROI 1. Fe^3+^ maps show that most of the Fe was in its oxidized form, with a mean of 82 % Fe^3+^ in ROI 1 and 66 % in ROI 2. Orange arrows in panel (A) indicate what appears to be the encrusted periplasm. The corresponding EDS and EELS spectra of these ROIs are presented in Figure S8.

Oxygen was the most abundant element in all biominerals, while the ratio of Fe atoms normalized to O atoms (Fe/O) ranged from 68 to 83% (mean 74 ± 5.3%) in the spherical NPs. In contrast, the crystalline “flakes” were richer in Fe, where the Fe/O varied from 81 to 93% (mean of 85 ± 5.3%). The atomic As to Fe ratio (normalized to Fe atoms) varied from 2.1 to 25% in all the regions analyzed (Table S1, Figures 5E & J and S9 F & L). “Flakes” showed lower atomic As/Fe ratios (2.1-6.4%) (Figures 5E & S9 F), while the cell-associated spherical NPs had higher atomic As/Fe ratios (11-25%) (Table S1 and Figures 5K & S9 L). Only one cell-detached NP was analyzed, which had 24% As/Fe (Figure S9 L). Thus, it appears that the crystalline “flakes” are low in oxygen and arsenic, in contrast to the spherical NPs, which retained more As (III/V), possibly due to a higher sorption capacity^57^. The atomic As/Fe/O ratios suggest that As-bearing minerals were not formed during these conditions, and As was retained through other processes, such as sorption or co-precipitation. Even though the “flakes” developed after a longer incubation period, and in principle were more thermodynamically stable^60^, this mineral phase was not the most efficient for As retention.

### 3.8 High-resolution Fe^3+^ mapping in biominerals using STEM and EELS

The Fe oxidation state in these biominerals was further mapped by STEM/EELS, where Fe^3+^ was predominant in both biomineral morphologies (Table S1 and Figures 5F & K and S9 G & M). Amorphous spherical NPs contained 66 to 98% Fe^3+^, while the crystalline biominerals contained 76 to 94% Fe^3+^. Although Fe^3+^ in Fe oxyhydroxides can undergo electron beam induced reduction^60, 61^, no electron beam damage was seen within these particles.

### 3.9 Scanning electron diffraction (SED) crystallinity maps

To obtain crystallinity maps scanning electron diffraction was used, which is a nanodiffraction technique. The diffraction pattern of the “flakes” showed bright spots, characteristic of single crystal specimens (inset in Figure 6C). The SED crystal identification phase map of a larger sample area identified lepidocrocite in the outer region of the sample (Figure 6C). This analysis agrees with the XRD results, indicating a correlative identification of the Fe(III)-oxyhydroxide lepidocrocite at the macro and nanoscale.

**Figure 6.**
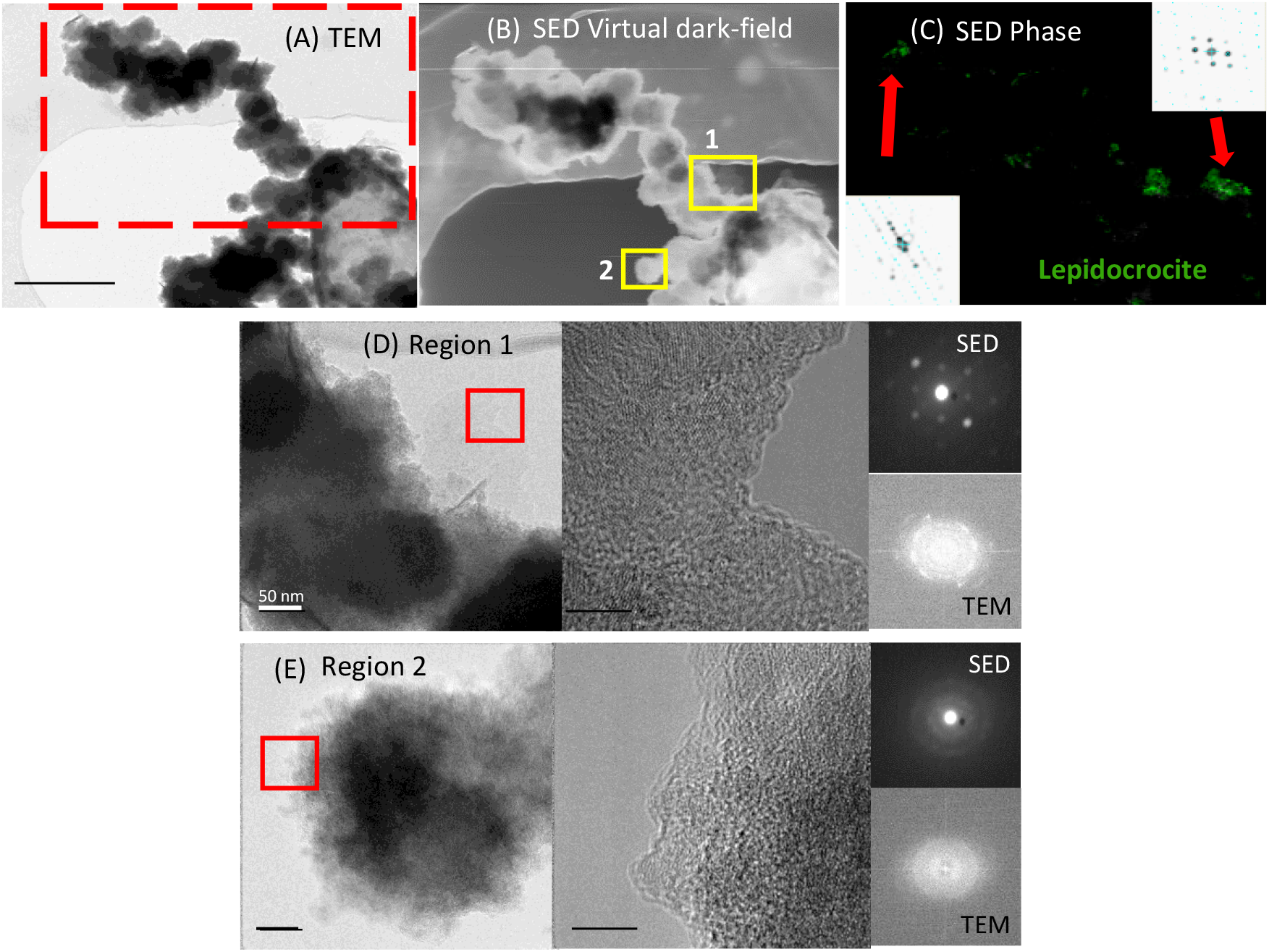
TEM and SED crystal identification maps from a mineralized ST3 cell pole and an extracellular fibril-like biomineral. (A) Bright-field TEM micrograph of the whole region analyzed (red dashed rectangle); (B) SED virtual dark-field micrograph of the region in the red dashed rectangle in panel (A); (C) SED crystal identification phase map of the whole region in panel (B), showing lepidocrocite in green, insets are the diffraction patterns, indicating crystalline structures; (D) Higher magnification TEM micrograph of Region 1 (yellow square #1 in (B)), with the HRTEM micrograph showing lattice fringes of the region selected in the red square in (D), and the corresponding SED and TEM diffraction patterns, indicating a crystalline structure; and (E) Higher magnification TEM micrograph of Region 2 (yellow square #2 in (B)), with the HRTEM micrograph showing lattice fringes of the region selected in the red square in (E), and the corresponding SED and TEM diffraction patterns, indicating an amorphous structure.

### 3.10 Biominerals produced by *Acidovorax* sp. strain ST3 and As sequestration

The aqueous phase analysis and biogenic minerals characterized here showed a high removal of As concomitant with Fe(III) precipitation in experiments with cells, confirming that cells facilitated Fe(II) oxidation and As precipitation/sorption. Functional groups on cells may influence the mineral formation and evolution, in contrast to extracellular mineral precipitation, where the medium composition may exert an additional effect. The Fe(III)-oxyhydroxide lepidocrocite [Fe^3+^O(OH)] was consistently identified with nanodiffraction in STEM after 7 days of incubation, and after 10 days both lepidocrocite and goethite were identified using XRD. Moreover, the expected oxidation state of Fe in lepidocrocite (Fe^3+^) was consistent with the Fe^3+^ identified by EELS. The Fe/O ratios determined by EDS contrast with these results, however, this estimation is sensible, considering that the quantification of light elements such as O is challenging in the presence of heavier elements ^62, 63^. Moreover, as absorption correction is not included in the STEM-EDS quantification, this analysis is likely to overestimate the Fe content, since higher energy Fe X-rays will be absorbed less strongly than the lower energy oxygen. Goethite and particularly lepidocrocite, can be susceptible to bioreduction^64, 65^, potentially remobilizing As. However, the crystalline “flakes” (developed after a longer incubation) were As deficient, in contrast to the amorphous spherical NPs, which retained more As, as shown by STEM-EDS analysis. Therefore, it appears that a longer incubation period would not necessarily promote higher As sequestration by these more crystalline biominerals in these conditions.

### 3.11 Identification of Fe encrustation sites in the cell to infer the Fe(II) oxidation mechanism

HAADF-STEM images are characterised by Z-contrast, where elements of high atomic number appear brighter^66^, for instance, the bright region in Figure 5A indicated by orange arrows, can be expected to be rich in Fe and As. This region matches the location and reported size of the periplasm (mean 25.8 ± 6.2 nm) using traditional TEM^67^. Therefore, these data are consistent with the periplasm being encrusted. Moreover, no cytoplasmic encrustation was observed. Although these micrographs are 2D projections of a 3D object, this periplasmic encrustation is consistent with the presented complementary NanoSIMS depth-profiles, where cells were probed by slices of a few tens of nm and remodeled in 3D (Figures 3 & S6).

The identification of the site of mineralization was used to infer the mechanism of Fe(II) oxidation in strain ST3. In contrast to periplasmic enzymatic As(III) oxidation^10^, the most plausible pathway for Fe(II) oxidation in strain ST3 is through reaction with nitrite, in agreement with other Fe(II)-oxidizers^16^, where Fe(II) is not only oxidized in the periplasm (due to periplasmic denitrification), but also extracellularly, as a consequence of reaction with the released nitrite species. Biomineralization patterns are characteristic of the specific Fe(II) oxidation mechanisms used by bacteria, and periplasmic biomineralization could be a common trait in the *Acidovorax* genus. In addition, hydrophilicity and surface charge are two factors that drive mineral precipitation at the cell surface, as reported for other Fe(II)-oxidizing bacteria^68^, and investigating these conditions warrants an advancement in understanding cell mineralization and metabolic heterogeneity in *Acidovorax* sp. strain ST3.

## 4. Conclusions

The NanoSIMS single cell analysis revealed metabolic heterogeneity with similarly heterogeneous mineralization patterns that suggest an interdependency between the two processes, most likely related to the growth stage of individual cells. Although periplasmic mineral encrustation disrupted metabolic activities, these mineralized cells retained As(III/V), in contrast to the active cells, that were practically devoid of this element.

The application of complementary nanoscale techniques was vital for a comprehensive analysis of Fe(II) and As(III) oxidation and mineralization under denitrifying conditions by *Acidovorax* sp. strain ST3. Bacteria facilitated Fe(II) oxidation to Fe(III), and although bacterial As(III) oxidation was not evident in the aqueous phase, As(III) removal was quantified in the aqueous phase in incubations with cells, and Fe(III) minerals sorbed or co-precipitated As(III/V) in varying ratios. NanoSIMS enabled simultaneous depth profiling of active/inactive and encrusted/non-encrusted cells with variable levels of metabolic activity, which could only be achieved with this technique. HAADF-STEM, EDS, SED and EELS allowed quantitative analysis of the composition of biominerals at high spatial resolution, and identified the intracellular compartment where cell encrustation occurs, complementary to the NanoSIMS 3D results. Arsenic was expected to spatially co-localize with Fe(III) biominerals in these conditions, which occurred in most cases. However, the NanoSIMS depth-profiles produced an insightful 3D analysis of the intracellular As and Fe distribution at high resolution and revealed unanticipated As and Fe spatial delocalization in some cells. This investigation advances our understanding of the molecular processes governing bacterial iron mineralization and its impact on trace metal fate.

## Author Contributions

RLA prepared the samples for aqueous and solid phase analysis, and performed the SEM, XRD and NanoSIMS data acquisition/analysis. JL and RLA designed the experiments. SMF performed the STEM/EDS/SED/EELS data acquisition/analysis. KLM and IL assisted in the NanoSIMS data acquisition/analysis. FJZ and JZ first isolated the ST3 strain and assisted with the initial experimental design. SJH assisted in STEM data interpretation. RLA wrote the main manuscript. All authors contributed to the article and approved the submitted version.

## Acknowledgements

RLA thanks the Mexican National Council for Science and Technology (CONACyT) for granting the PhD scholarship number 411911. We thank the support of the Natural Environment Research Council (GOAM NE/P01304X/1). Thanks to Mr Paul Lythgoe and Mr Alastair Bewsher for performing aqueous geochemistry analysis. We thank Dr John Waters for his help in XRD spectra interpretation and Dr Kexue Li for the training in Avizo^®^ NanoSIMS 3D reconstructions. The NanoSIMS was funded by UK Research Partnership Investment Funding (UKRPIF) Manchester RPIF Round 2. This work was supported by the Henry Royce Institute for Advanced Materials, funded through EPSRC grants EP/R00661X/1, EP/S019367/1, EP/P025021/1 and EP/P025498/1, and National Natural Science Foundation of China (41930758). S.J.H and S.M.F. acknowledge funding from the European Research Council (Horizon 2020, grant agreement ERC-2016-STG-EvoluTEM-715502) and the Rosalind Franklin Institute’s Correlated Imaging theme, funded by EPSRC grant(s) EP/S001999/1 and EP/T006889/1.

## References

1. R. J. Bowell, C. N. Alpers, H. E. Jamieson, D. K. Nordstrom and J. Majzlan, The Environmental Geochemistry of Arsenic -- An Overview, Reviews in Mineralogy and Geochemistry, 2014, 79, 1–16.

2. D. E. Cummings, F. Caccavo, S. Fendorf and R. F. Rosenzweig, Arsenic Mobilization by the Dissimilatory Fe(III)-Reducing Bacterium Shewanella alga BrY, Environmental Science and Technology, 1999, 33, 723–729.

3. F. Islam, A. G. Gault, C. Boothman, D. A. Polya, J. M. Charnock, D. Chatterjee and J. R. Lloyd, Role of metal-reducing bacteria in arsenic release from Bengal delta sediments, Nature, 2004, 430, 68–71.

4. J. R. Lloyd and R. S. Oremland, Microbial transformations of Arsenic in the environment-from Soda lakes to Aquifers, Elements, 2006, 2, 85–90.

5. J.-H. Huang, Impact of Microorganisms on Arsenic Biogeochemistry: A Review, Water, Air, & Soil Pollution, 2014, 225.

6. K. M. Campbell and D. K. Nordstrom, Arsenic Speciation and Sorption in Natural Environments, Reviews in Mineralogy and Geochemistry, 2014, 79, 185–216.

7. J. P. Shapleigh, in The Prokaryotes. A Handbook on the Biology of Bacteria, eds. M. Dworkin, S. Falkow, E. Rosenberg, K.-H. Schleifer and E. Stackebrandt, Springer, USA, 2006, DOI: 10.1007/0-387-30742-7_23, ch. Chapter 23, pp. 769–792.

8. E. D. Rhine, C. D. Phelps and L. Y. Young, Anaerobic arsenite oxidation by novel denitrifying isolates, Environ Microbiol, 2006, 8, 899–908.

9. J. Zhang, W. Zhou, B. Liu, J. He, Q. Shen and F. J. Zhao, Anaerobic arsenite oxidation by an autotrophic arsenite-oxidizing bacterium from an arsenic-contaminated paddy soil, Environ Sci Technol, 2015, 49, 5956–5964.

10. J. Zhang, S. Zhao, Y. Xu, W. Zhou, K. Huang, Z. Tang and F. J. Zhao, Nitrate Stimulates Anaerobic Microbial Arsenite Oxidation in Paddy Soils, Environ Sci Technol, 2017, 51, 4377–4386.

11. Y. G. Zhu, M. Yoshinaga, F. J. Zhao and B. P. Rosen, Earth Abides Arsenic Biotransformations, Annu Rev Earth Planet Sci, 2014, 42, 443–467.

12. H. K. Carlson, I. C. Clark, S. J. Blazewicz, A. T. Iavarone and J. D. Coates, Fe(II) oxidation is an innate capability of nitrate-reducing bacteria that involves abiotic and biotic reactions, Journal of Bacteoriology, 2013, 195, 3260–3268.

13. A. Price, M. C. Macey, J. Miot and K. Olsson-Francis, Draft Genome Sequences of the Nitrate-Dependent Iron-Oxidizing Proteobacteria Acidovorax sp. Strain BoFeN1 and Paracoccus pantotrophus Strain KS1, Microbiol Resour Announc, 2018, 7.

14. A. Kappler, C. Bryce, M. Mansor, U. Lueder, J. M. Byrne and E. D. Swanner, An evolving view on biogeochemical cycling of iron, Nature reviews. Microbiology, 2021, 19, 360–374.

15. A. Kappler, B. Schink and D. K. Newman, Fe(III) mineral formation and cell encrustation by the nitrate-dependent Fe(II)-oxidiser strain BoFeN1, Geobiology, 2005, 3, 235–245.

16. J. Zhang, C. W. Chai, L. K. ThomasArrigo, S. C. Zhao, R. Kretzschmar and F. J. Zhao, Nitrite Accumulation Is Required for Microbial Anaerobic Iron Oxidation, but Not for Arsenite Oxidation, in Two Heterotrophic Denitrifiers, Environ Sci Technol, 2020, 54, 4036–4045.

17. N. Klueglein and A. Kappler, Abiotic oxidation of Fe(II) by reactive nitrogen species in cultures of the nitrate-reducing Fe(II) oxidizer Acidovorax sp. BoFeN1 - questioning the existence of enzymatic Fe(II) oxidation, Geobiology, 2013, 11, 180–190.

18. J. Jamieson, H. Prommer, A. H. Kaksonen, J. Sun, A. J. Siade, A. Yusov and B. Bostick, Identifying and Quantifying the Intermediate Processes during Nitrate-Dependent Iron(II) Oxidation, Environ Sci Technol, 2018, 52, 5771–5781.

19. S. K. Chaudhuri, J. G. Lack and J. D. Coates, Biogenic magnetite formation through anaerobic biooxidation of Fe(II), Applied and environmental microbiology, 2001, 67, 2844–2848.

20. J. Sun, S. N. Chillrud, B. J. Mailloux, M. Stute, R. Singh, H. Dong, C. J. Lepre and B. C. Bostick, Enhanced and stabilized arsenic retention in microcosms through the microbial oxidation of ferrous iron by nitrate, Chemosphere, 2016, 144, 1106–1115.

21. J. Miot, J. Li, K. Benzerara, M. T. Sougrati, G. Ona-Nguema, S. Bernard, J.-C. Jumas and F. Guyot, Formation of single domain magnetite by green rust oxidation promoted by microbial anaerobic nitrate-dependent iron oxidation, Geochimica et Cosmochimica Acta, 2014, 139, 327–343.

22. L. Zhao, H. Dong, R. Kukkadapu, A. Agrawal, D. Liu, J. Zhang and R. E. Edelmann, Biological oxidation of Fe(II) in reduced nontronite coupled with nitrate reduction by Pseudogulbenkiania sp. Strain 2002, Geochimica et Cosmochimica Acta, 2013, 119, 231–247.

23. C. Hohmann, E. Winkler, G. Morin and A. Kappler, Anaerobic Fe(II)-oxidising bacteria show As resistance and immobilize As during Fe(III) mineral precipitation, Environmental science & technology, 2010, 44, 94–101.

24. P. Larese-Casanova, S. B. Haderlein and A. Kappler, Biomineralization of lepidocrocite and goethite by nitrate-reducing Fe(II)-oxidizing bacteria: Effect of pH, bicarbonate, phosphate, and humic acids, Geochimica et Cosmochimica Acta, 2010, 74, 3721–3734.

25. A. Kappler and D. K. Newman, Formation of Fe(III)-minerals by Fe(II)-oxidizing photoautotrophic bacteria, Geochimica et Cosmochimica Acta, 2004, 68, 1217–1226.

26. W. Xiu, H. Guo, J. Shen, S. Liu, S. Ding, W. Hou, J. Ma and H. Dong, Stimulation of Fe(II) Oxidation, Biogenic Lepidocrocite Formation, and Arsenic Immobilization by Pseudogulbenkiania Sp. Strain 2002, Environ Sci Technol, 2016, 50, 6449–6458.

27. S. Li, X. Li and F. Li, Fe(II) oxidation and nitrate reduction by a denitrifying bacterium, Pseudomonas stutzeri LS-2, isolated from paddy soil, Journal of Soils and Sediments, 2017, 18, 1668–1678.

28. K. Benzerara, G. Morin, T. H. Yoon, J. Miot, T. Tyliszczak, C. Casiot, O. Bruneel, F. Farges and G. E. Brown, Nanoscale study of As biomineralization in an acid mine drainage system, Geochimica et Cosmochimica Acta, 2008, 72, 3949–3963.

29. G. Schmid, F. Zeitvogel, L. Hao, P. Ingino, M. Floetenmeyer, Y. D. Stierhof, B. Schroeppel, C. J. Burkhardt, A. Kappler and M. Obst, 3-D analysis of bacterial cell-(iron)mineral aggregates formed during Fe(II) oxidation by the nitrate-reducing Acidovorax sp. strain BoFeN1 using complementary microscopy tomography approaches, Geobiology, 2014, 12, 340–361.

30. T. Suzuki, H. Hashimoto, N. Matsumoto, M. Furutani, H. Kunoh and J. Takada, Nanometer-scale visualization and structural analysis of the inorganic/organic hybrid structure of Gallionella ferruginea twisted stalks, Applied and environmental microbiology, 2011, 77, 2877–2881.

31. J. Baumgartner, G. Morin, N. Menguy, T. Perez Gonzalez, M. Widdrat, J. Cosmidis and D. Faivre, Magnetotactic bacteria form magnetite from a phosphate-rich ferric hydroxide via nanometric ferric (oxyhydr)oxide intermediates, Proc Natl Acad Sci U S A, 2013, 110, 14883–14888.

32. J. Miot, K. Benzerara, G. Morin, A. Kappler, S. Bernard, M. Obst, C. Férard, F. Skouri-Panet, J.-M. Guigner, N. Posth, M. Galvez, G. E. Brown and F. Guyot, Iron biomineralization by anaerobic neutrophilic iron-oxidizing bacteria, Geochimica et Cosmochimica Acta, 2009, 73, 696–711.

33. J. Miot, S. Lu, G. Morin, A. Adra, K. Benzerara and K. Küsel, Iron mineralogy across the oxycline of a lignite mine lake, Chemical Geology, 2016, 434, 28–42.

34. J. Miot, D. Jézéquel, K. Benzerara, L. Cordier, S. Rivas-Lamelo, F. Skouri-Panet, C. Férard, M. Poinsot and E. Duprat, Mineralogical Diversity in Lake Pavin: Connections with Water Column Chemistry and Biomineralization Processes, Minerals, 2016, 6, 24.

35. J. Miot, L. Remusat, E. Duprat, A. Gonzalez, S. Pont and M. Poinsot, Fe biomineralization mirrors individual metabolic activity in a nitrate-dependent Fe(II)-oxidizer, Front Microbiol, 2015, 6, 879.

36. C. Tominski, T. Lösekann-Behrens, A. Ruecker, N. Hagemann, S. Kleindienst, C. W. Mueller, C. Höschen, I. Kögel-knabner, A. Kappler and S. Behrens, Insights into carbon metabolisms provided by FISH-SIMS imaging of an autotrophic, nitrate reducing Fe(II) oxidizing enrichment culture, Appl. Envir. Microbiol., 2018, 84, 1–19.

37. M. Ilbert and V. Bonnefoy, Insight into the evolution of the iron oxidation pathways, Biochim Biophys Acta, 2013, 1827, 161–175.

38. A. Chakraborty, E. E. Roden, J. Schieber and F. Picardal, Enhanced growth of Acidovorax sp. strain 2AN during nitrate-dependent Fe(II) oxidation in batch and continuous-flow systems, Applied and environmental microbiology, 2011, 77, 8548–8556.

39. J. Miot, K. Benzerara, M. Obst, A. Kappler, F. Hegler, S. Schadler, C. Bouchez, F. Guyot and G. Morin, Extracellular iron biomineralization by photoautotrophic iron-oxidizing bacteria, Applied and environmental microbiology, 2009, 75, 5586–5591.

40. S. Park, D. H. Kim, J. H. Lee and H. G. Hur, Sphaerotilus natans encrusted with nanoball-shaped Fe(III) oxide minerals formed by nitrate-reducing mixotrophic Fe(II) oxidation, FEMS Microbiol Ecol, 2014, 90, 68–77.

41. M. Nordhoff, C. Tominski, M. Halama, J. M. Byrne, M. Obst, S. Kleindienst, S. Behrens and A. Kappler, Insights into Nitrate-Reducing Fe(II) Oxidation Mechanisms through Analysis of Cell-Mineral Associations, Cell Encrustation, and Mineralogy in the Chemolithoautotrophic Enrichment Culture KS, Applied and Environmental Microbiology, 2017, 83, 1–19.

42. B. Li, X. Pan, D. Zhang, D.-J. Lee, F. A. Al-Misned and M. G. Mortuza, Anaerobic nitrate reduction with oxidation of Fe(II) by Citrobacter Freundii strain PXL1 – a potential candidate for simultaneous removal of As and nitrate from groundwater, Ecological Engineering, 2015, 77, 196–201.

43. J. Miot, K. Benzerara, G. Morin, S. Bernard, O. Beyssac, E. Larquet, A. Kappler and F. Guyot, Transformation of vivianite by anaerobic nitrate-reducing iron-oxidizing bacteria, Geobiology, 2009, 7, 373–384.

44. F. Hegler, N. R. Posth, J. Jiang and A. Kappler, Physiology of phototrophic iron(II)-oxidizing bacteria: implications for modern and ancient environments, FEMS Microbiol Ecol, 2008, 66, 250–260.

45. D. R. Lovley and E. J. P. Phillips, Organic matter mineralization with reduction of ferric iron in anaerobic sediments, Applied and Environmental Microbiology, 1986, 51, 683–689.

46. A. G. Gault, J. Jana, S. Chakraborty, P. Mukherjee, M. Sarkar, B. Nath, D. A. Polya and D. Chatterjee, Preservation strategies for inorganic arsenic species in high iron, low-Eh groundwater from West Bengal, India, Anal Bioanal Chem, 2005, 381, 347–353.

47. D. McPhail and M. Dowsett, in Surface Analysis.The Principal Techniques, eds. J. C. Vickerman and I. S. Gilmore, John Wiley & Sons, United Kingdom, 2nd edn., 2009, ch. 5, pp. 207–268.

48. R. Lopez-Adams, L. Newsome, K. L. Moore, I. C. Lyon and J. R. Lloyd, Dissimilatory Fe(III) Reduction Controls on Arsenic Mobilization: A Combined Biogeochemical and NanoSIMS Imaging Approach, Front Microbiol, 2021, 12, 640734.

49. P. A. van Aken and B. Liebscher, Quantification of ferrous/ferric ratios in minerals: new evaluation schemes of Fe L 23 electron energy-loss near-edge spectra, Physics and Chemistry of Minerals, 2002, 29, 188–200.

50. J. Gorny, G. Billon, L. Lesven, D. Dumoulin, B. Made and C. Noiriel, Arsenic behavior in river sediments under redox gradient: a review, Sci Total Environ, 2015, 505, 423–434.

51. C. Bryce, N. Blackwell, C. Schmidt, J. Otte, Y. M. Huang, S. Kleindienst, E. Tomaszewski, M. Schad, V. Warter, C. Peng, J. M. Byrne and A. Kappler, Microbial anaerobic Fe(II) oxidation -Ecology, mechanisms and environmental implications, Environ Microbiol, 2018, 20, 3462–3483.

52. S. French, D. Puddephatt, M. Habash and S. Glasauer, The dynamic nature of bacterial surfaces: implications for metal-membrane interaction, Crit Rev Microbiol, 2013, 39, 196–217.

53. S. Shen, X. F. Li, W. R. Cullen, M. Weinfeld and X. C. Le, Arsenic binding to proteins, Chem Rev, 2013, 113, 7769–7792.

54. L. J. Bird, M. L. Coleman and D. K. Newman, Iron and copper act synergistically to delay anaerobic growth of bacteria, Applied and environmental microbiology, 2013, 79, 3619–3627.

55. V. Takhaveev and M. Heinemann, Metabolic heterogeneity in clonal microbial populations, Curr Opin Microbiol, 2018, 45, 30–38.

56. E. Simsek and M. Kim, The emergence of metabolic heterogeneity and diverse growth responses in isogenic bacterial cells, ISME J, 2018, 12, 1199–1209.

57. S. Dixit and J. G. Hering, Comparison of Arsenic(V) and Arsenic(III) Sorption onto Iron Oxide Minerals: Implications for Arsenic Mobility, Environ Sci Technol, 2003, 37, 4182–4189.

58. D. Mavrocordatos and D. Fortin, Quantitative characterization of biotic iron oxides by analytical electron microscopy, American Mineralogist, 2002, 87, 940–946.

59. M. Sanchez-Roman, F. Puente-Sanchez, V. Parro and R. Amils, Nucleation of Fe-rich phosphates and carbonates on microbial cells and exopolymeric substances, Front Microbiol, 2015, 6, 1024.

60. A. Gloter, M. Zbinden, F. Guyot, F. Gaill and C. Colliex, TEM-EELS study of natural ferrihydrite from geological–biological interactions in hydrothermal systems, Earth and Planetary Science Letters, 2004, 222, 947–957.

61. Y.-H. Pan, G. Vaughan, R. Brydson, A. Bleloch, M. Gass, K. Sader and A. Brown, Electron-beam-induced reduction of Fe3+ in iron phosphate dihydrate, ferrihydrite, haemosiderin and ferritin as revealed by electron energy-loss spectroscopy, Ultramicroscopy, 2010, 110, 1020–1032.

62. A. Warley, Development and comparison of the methods for quantitative electron probe X-ray microanalysis analysis of thin specimens and their application to biological material, J Microsc, 2016, 261, 177–184.

63. G. M. Roomans, Introduction to X-Ray Microanalysis in Biology, Journal of Electron Microscopy Technique, 1988, 9, 3–17.

64. E. J. O’Loughlin, P. Larese-Casanova, M. Scherer and R. Cook, Green Rust Formation from the Bioreduction of γ –FeOOH (Lepidocrocite): Comparison of SeveralShewanellaSpecies, Geomicrobiology Journal, 2007, 24, 211–230.

65. Y. Dong, R. A. Sanford, M. I. Boyanov, T. M. Flynn, E. J. O’Loughlin, K. M. Kemner, S. George, K. E. Fouke, S. Li, D. Huang, S. Li and B. W. Fouke, Controls on Iron Reduction and Biomineralization over Broad Environmental Conditions as Suggested by the Firmicutes Orenia metallireducens Strain Z6, Environ Sci Technol, 2020, 54, 10128–10140.

66. R. F. Egerton, Physical principles of electron microscopy: an introduction to TEM, SEM and AEM, Springer, USA, 2005.

67. J. Miot, K. Maclellan, K. Benzerara and N. Boisset, Preservation of protein globules and peptidoglycan in the mineralized cell wall of nitrate-reducing, iron(II)-oxidizing bacteria: a cryo-electron microscopy study, Geobiology, 2011, 9, 459–470.

68. G. Saini and C. S. Chan, Near-neutral surface charge and hydrophilicity prevent mineral encrustation of Fe-oxidizing micro-organisms, Geobiology, 2013, 11, 191–200.

