## Supplementary data file for "Elucidating Heterogeneous Iron Biomineralization Patterns in a Denitrifying As(III)-Oxidizing Bacterium: Implications for Arsenic Immobilization"

**Electronic Supplementary information**

**for**

### Text S1. Supplementary materials and methods

#### Text S1.1 Biofilm sample preservation for high vacuum analysis (SEM and NanoSIMS)

The Si wafers from biofilm experiments were removed from the bottles in an anaerobic chamber at days 7 and 14 and preserved by chemical fixation with glutaraldehyde and dehydration with ethanol, following a procedure detailed elsewhere[1]. The wafers were removed from the last ethanol solution and air-dried in the anaerobic chamber overnight. Wafers collected at day 7 were also preserved by only rinsing with $_{d}H_2O$ and air-dried in the anaerobic chamber, to compare the cell and mineral preservation with this approach, used in TEM sample preparation. Once dried overnight, all the Si wafers were sputter coated with 10 nm of platinum and kept in anoxic conditions until analysis.

#### Text S1.2 Planktonic sample preparation

Planktonic samples were collected at days 4 and 7 for STEM and on days 7 and 10 for XRD analysis. Samples (1 mL) were centrifuged (2509 g/30 min) and washed twice in $N_2$-degassed deionised water ($_{d}H_2O$). For STEM samples the washed precipitates were re-suspended in $_{d}H_2O$ to an $OD_{600}$≈0.1, this dilution (1.5 µL) was placed onto lacey carbon films on Cu TEM grids (Agar Scientific®) and further dehydrated using a turbo pump. For XRD analysis the concentrated precipitates (2 mL) were spread on glass slides (1 x 0.5 cm) and dried overnight in an anoxic cabinet All these steps, including sample handling and loading into the instruments, were performed under anoxic conditions.

#### Text S1.3 STEM anoxic sample loading

Sample transportation to the TEM laboratory was done inside an airtight container in an anoxically sealed plastic bag. The TEM grids loading onto the instrument sample holder was done using a plastic box (~70 x 40 x 40 cm) filled with argon at the bottom layer, and the transfer of the TEM grid from the portable sample holder to the TEM instrument sample holder was done below the height of the argon layer filling inside the plastic box. This sample manipulation was challenging but with the aim to keep the anoxic conditions for as long as possible, although the presence of some ppm of $O_2$ exposure during specimen loading cannot be ruled out.

#### Text S1.4 NanoSIMS analysis details

In NanoSIMS analysis the elemental ratio $^{13}C/^{12}C$ was used to identify cells that were metabolically active; $^{13}C^{14}N/^{12}C^{14}N$ was not chosen due to the unresolvable isobaric interferences $^{10}B^{16}O$ and

$^{11}B^{16}O$. The molecular ions $^{56}Fe^{12}C^-$ and $^{56}Fe^{16}O^-$ were used to monitor $^{56}Fe$ because $^{56}Fe^-$ has a low ionisation yield under the $Cs^+$ ion beam [2].

**Text S1.5 NanoSIMS depth profiles**

Single cells were depth profiled using both primary ion beams in NanoSIMS. Isolated cells on the Si wafer were selected for this, and these cells were not implanted to reach steady state, to keep the cell mineralization coating intact. However, a quick implantation of $Cs^+$ or $O^-$ ions (D1=1, 30 s) was done before the start of analysis to locate and focus single cells. In NanoSIMS, $^{56}Fe$ has a higher secondary ion yield under $O^-$ bombardment than under $Cs^+$ ions bombardment [2], where $^{56}Fe$ is normally collected as the molecular ion $^{56}Fe^{16}O^-$ which enhances detection. Therefore, $^{56}Fe^+$ produced higher count intensities, typically one order of magnitude higher than $^{56}Fe^{16}O^-$ (Figures 3C, 3H, S5 C & S5 H). This is the opposite for $^{75}As$, because $^{75}As$ has a lower ion yield under the $O^-$ beam than under $Cs^+$ bombardment (Figures 3D, 3I, S5 D & S5 I). In these samples the $Cs^+$ beam was useful to infer metabolic activity by $^{13}C$ accumulation and to map As and Fe, while the $O^-$ beam was particularly useful to map $Fe^+$, because it produced a stronger signal.

**Text S1.5.1 NanoSIMS depth profiles using $Cs^+$ primary ions**

During $Cs^+$ depth profiling D1 apertures were reduced to 4 or 5, reducing the total current. Pixel sizes were 128 x 128 or 256 x 256, and dwell time varied from 5,000-20,000 µs $px^{-1}$. The number of planes collected were in the range of 50-170, and the scanning was stopped when the $^{12}C$ or $^{12}C^{14}N$ signal disappeared, indicating that the bacterial cell had been entirely sputtered away and chemical information could be recovered from the whole cell.

**Text S1.5.2 NanoSIMS depth profiles using $O^-$ primary ions**

The $O^-$ beam was scanned on the sample surface with a current between 1.693-4.31 pA (D1=4-5). Images were collected at a dwell time of 5000 µs $px^{-1}$. The positive secondary ions simultaneously detected were: $^{23}Na$, $^{24}Mg$, $^{28}Si$, $^{39}K$, $^{44}Ca$, $^{56}Fe$ and $^{75}As$. CAMECA MRP was improved to ≈3000 (ES=3 AS=2) to separate the mass interferences $^{12}C^{16}O^+$ at mass 28 and $^{23}Na^{16}O^+$ at mass 39. Mass 75 had no isobaric interferences ($^{56}Fe^{19}F^+$ was not formed). In this case the disappearance of the $^{56}Fe^+$ signal was monitored as an indicator of complete cell sputtering. The ionic salts (Na, Mg, K and Ca) were only used as a reference of cell sputtering and their mobility during the chemical fixation process is acknowledged [3, 4].

#### Text S1.5.3 NanoSIMS data analysis

L'image software was used to obtain stack NanoSIMS images (Larry Nittler, Carnegie Institution of Washington). ImageJ (https://imagej.nih.gov/ij/) with the plugin OpenMIMS (MIMS, Harvard University; www.nrims.harvard.edu) was used to create the hue saturation images (HSI) isotope ratio maps and to generate colour merge (overlay) images. Regions of interest (ROIs) were manually drawn around cells, the ion counts were normalized to primary ion doses.

#### Text S1.5.4 NanoSIMS single-cell 3D reconstructions

3D reconstructions of the depth profiles were created using the Thermo Scientific™ Avizo™ Software 9.7.0. Stack data of the negative secondary ions $^{12}C^{-}$, $^{56}Fe^{16}O^{-}$ and $^{75}As^{-}$ , and the positive secondary ions $^{23}Na^{+}$, $^{56}Fe^{+}$, and $^{75}As^{+}$, were first extracted with ImageJ and saved in the ".raw" format file. Afterwards, these .raw files were loaded into Avizo™. The Z depth was compressed to 15-20 % to reduce the space between planes (generated by the software), bringing the volume of 3D visualisation to scale. The $^{12}C^{-}$ and $^{23}Na^{+}$ signals were used to generate the bacterial surface by smoothing (averaging) the signal over 2-3 pixels. The commands "generate surface" and "show surface" were used sequentially, and the "transparent" display with a transparency of 80 % was selected. $^{56}Fe^{+}$ and $^{56}Fe^{16}O^{-}$ signals were the proxies of cellular Fe encrustation, smoothed to 2 pixels and displayed as "shaded". The $^{75}As^{+/-}$ ion counts were smoothed to 1 pixel, because of the lower ion counts, and displayed as "points" to enable their visualisation.

### Text S2. Supplementary results

#### Text S2.1 Sample preservation effect imaged in SEM

Sample preservation is often overlooked but it is a key determining factor for accurate image analyses, including electron microscopy and SIMS techniques. For this reason, two preservation techniques were used (air-drying and chemical fixation-dehydration) for the biofilm samples after 4 and 7 days of incubation, and these were imaged in SEM to compare the effect on cells and biominerals. Abundant biomass colonising the Si wafer was observed in the samples by day 7, regardless of the preparation method used (Figure S4 A & E), suggesting that both methods preserved biomass density, although substantial surface colonization was already present by day 4 (Figure S4 A). The chemical fixation-dehydration approach preserved the structure of the cells (Figure S4 F), in contrast to the air-dried samples, which showed many disrupted cells (Figure S3 B). Heavy mineralization of cells was observed after 7 days of incubation in both sample treatments, however, the cells in the chemically fixed samples showed biominerals in the shape of sheaths (yellow arrows in Figure S4 F), while the air-dried samples showed nanoparticles of a few tens of nm on the cell surfaces (magenta arrows in Figure S4 D). The Si wafer surfaces were coated with small particles, and these solids were similar in morphology to the biominerals on the surface of the cells with each type of sample preparation (Figure S4 D & F). Moreover, air-dried samples showed a residue covering the cells (orange arrows in Figure S4 C), possibly extracellular polysaccharides substances (EPS) or other organic material, which was not observed in the chemically fixed-dehydrated samples. For this reason and for the better preservation of cells, chemical fixation-dehydration was selected for NanoSIMS samples. However, the air-drying approach introduced less chemicals and possibly less artifacts that could alter the biomineral morphology, thus, the samples for biomineral characterization were treated by this method.

### S3. Supplementary tables

**Table S1**. Stoichiometry of As and O atoms normalized to Fe atoms in selected regions of interest (ROI) of mineralized cells or extracellular biominerals, obtained from the STEM-EDS or EELS maps, as indicated. Values are averaged over the whole ROI and expressed as percentage (%). For EDS only the elements As, Fe and O are quantified so these necessarily sum to 100%. For EELS only the ratio of $Fe^{3+}/Fe^{2+}$ is quantified.

| | | EDS | | | | | EELS |
|---|---|---|---|---|---|---|---|
| **ROI[a]** | **Morphology** | **As** | **Fe** | **O** | **As/Fe** | **Fe/O** | **$Fe^{3+}$** |
| 1 (ROI 1 Fig. 5) | Amorphous | 8.9 | 37.6 | 53.4 | 24 | 70 | 66 |
| 2 (ROI 2 Fig. 5) | Crystalline | 1.1 | 47.6 | 51.2 | 2.4 | 93 | 82 |
| 3 (ROI 1 Fig. S8) | Crystalline | 2.8 | 43.4 | 53.8 | 6.4 | 81 | 76 |
| 4 (ROI 2 Fig. S8) | Amorphous | 9.4 | 39.2 | 51.4 | 24 | 77 | 72 |
| 5 | Amorphous | 4.3 | 39 | 56.6 | 11 | 68 | 98 |
| 6 | Amorphous | 9.8 | 38.5 | 51.7 | 25 | 75 | 69 |
| 7 | Amorphous | 9 | 38.4 | 52.6 | 23 | 73 | N/A |
| 8 | Crystalline | 1.1 | 44.4 | 54.4 | 2.5 | 82 | 94 |
| 9 | Crystalline | 1.7 | 44.6 | 53.7 | 3.8 | 82 | 88 |
| 10 | Crystalline | 1 | 46.7 | 52.2 | 2.1 | 89 | 88 |
| 11 | Amorphous | 5.7 | 42.5 | 51.8 | 13.5 | 83 | N/A |

[a]The first column shows in parenthesis the Figures in which the ROIs micrographs are shown where applicable.

[b]N/A= data not available.

### S4. Supplementary Figures

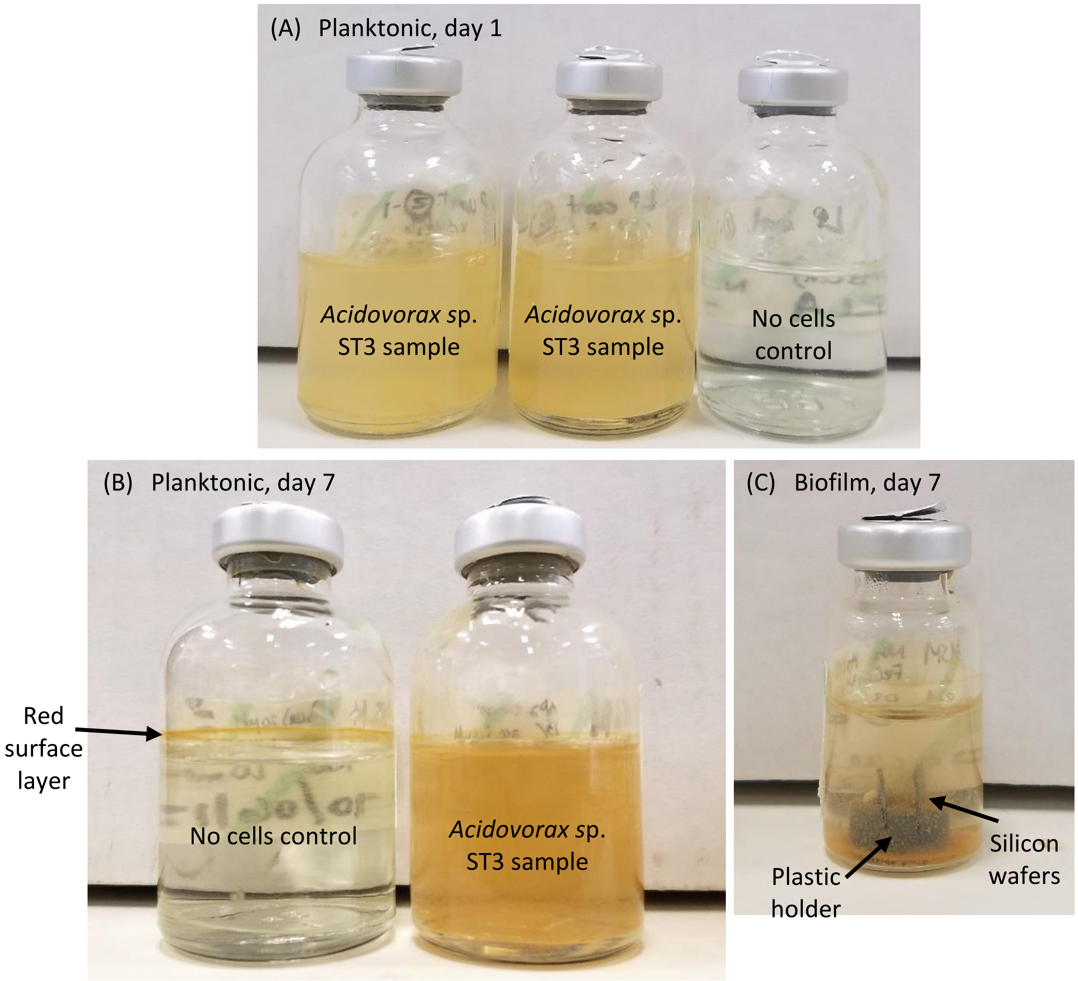


**Figure S1**. Pictures of the experimental bottles with *Acidovorax* sp. strain ST3 cells after 1 and 7 days of incubation. (A) planktonic growth samples at day 1, (B) planktonic growth samples at day 7 and (C) biofilm growth sample at day 7. In (B) the bottle on the left is the no cells control, showing the thin red layer that formed abiotically at the surface in some bottles.

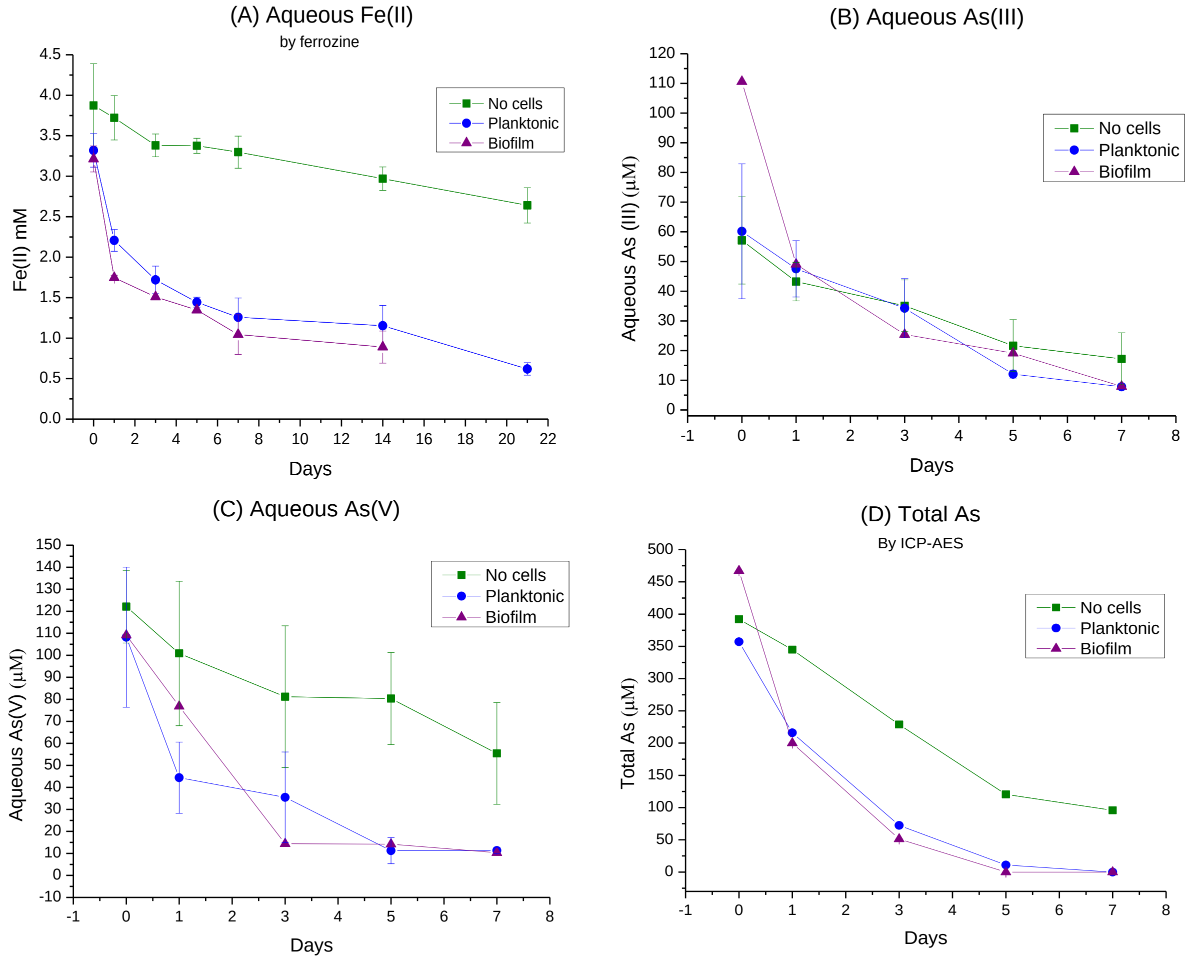


**Figure S2.** Fe(II) and As species measured in the aqueous phase in planktonic and biofilm cultures of strain ST3. (A) Fe(II) by ferrozine with planktonic and the no cell control measured to 21 days of incubation and biofilm growth measured to 14 days; (B-D) As species were all measured until 7 days of incubation: (B) As(III), (C) As(V) and (D) total As by ICP-AES. Notice the abiotic removal of As from solution, probably through sorption to abiotically formed precipitates or minerals. Error bars are the standard deviation, N=3.

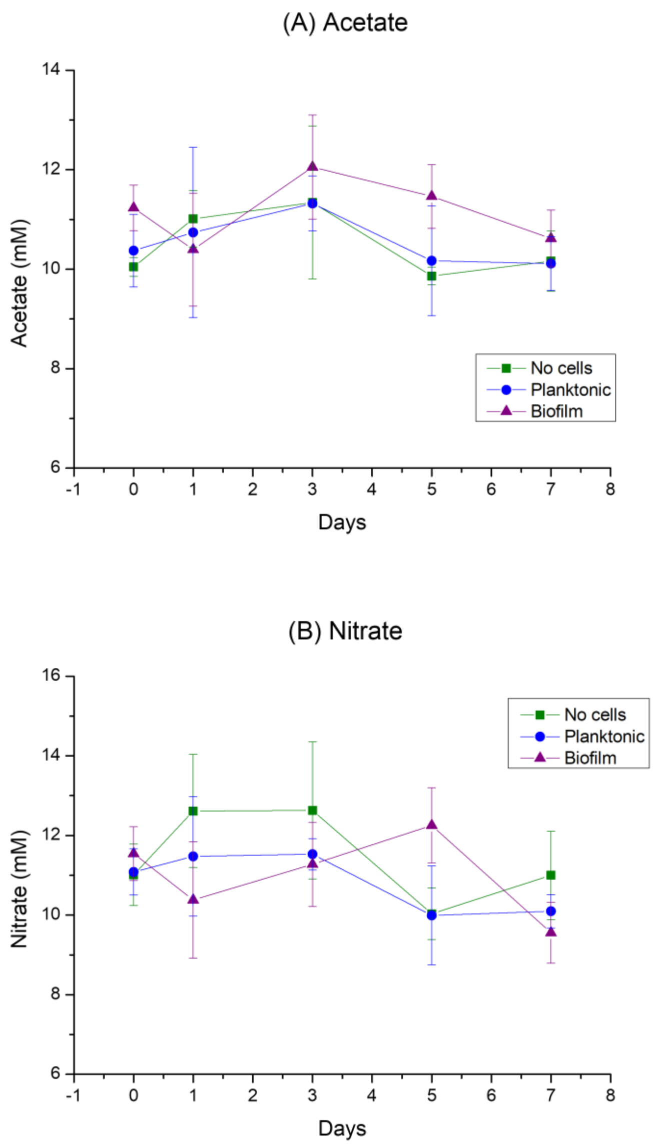


**Figure S3.** Acetate and nitrate quantification in the aqueous phase of strain ST3 grown in biofilm and planktonic conditions (as well as no cell control). Error bars are the standard deviation, N=3.

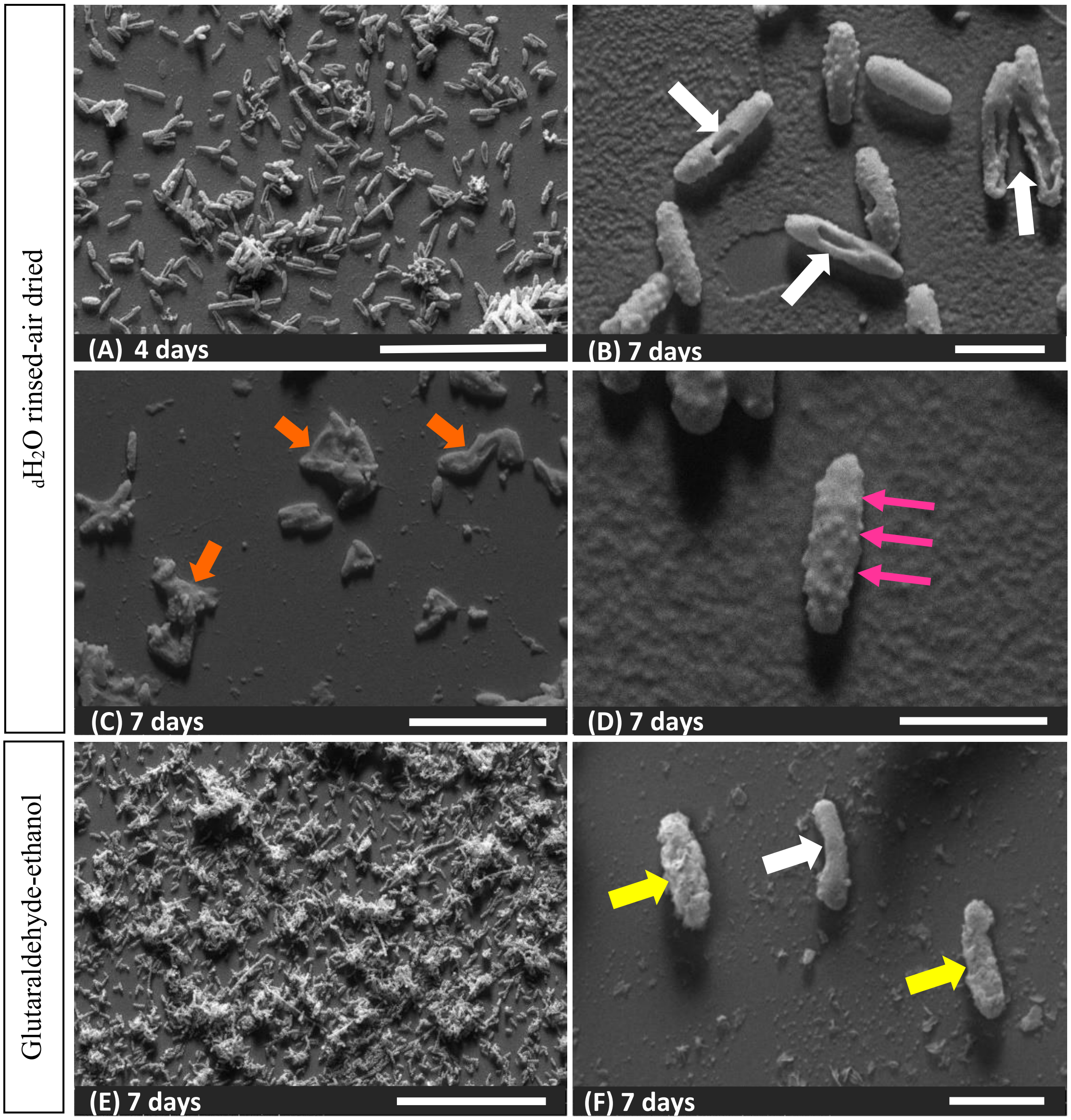


**Figure S4.** Contrasting sample preservation methods by SEM imaging of *Acidovorax* sp. ST3 cells grown in biofilm conditions. (A) $_{d}H_2O$ rinsed-air dried cells after 4 days and (B, C & D) 7 days of incubation, (E & F) are cells preserved by chemical fixation-dehydration after 7 days of incubation. Abundant biomass is observed in both preservation methods from day 4 of incubation (only shown for $_{d}H_2O$ rinsed-air dried cells in A), but differences are noticeable at the cell level and the morphology of the biominerals on the cell surface. The white arrow in (F) indicates a well-preserved cell with low mineralization while the white arrows in (B) indicate disrupted cells. The yellow arrows in (F) indicate heavily mineralized cells showing "flakes" of biominerals on the cell surface. The pink arrows in (D) show spherical particles of tens of nm on the cell surface. The orange arrows in (C) indicate groups of cells covered by a residue (potentially EPS). Scale bars are 20, 50 and 10 µm in (A), (C) and (E) respectively, and 2 µm in (B), (D) and (F).

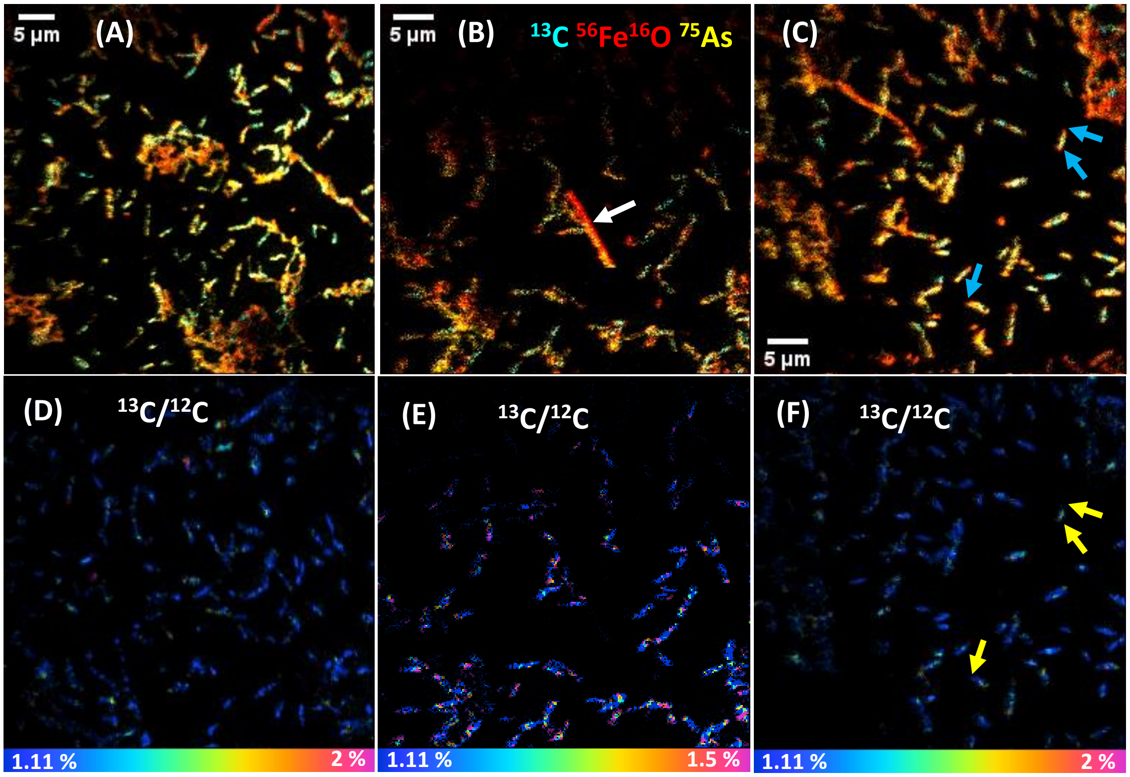


**Figure S5.** NanoSIMS images of strain ST3 biofilm growth at day 7. Top row (A-C) are overlay images of cells displaying $^{13}C$ (cyan), $^{56}Fe^{16}O$ (red) and $^{75}As$ (yellow). Bottom row (D-F) are the $^{13}C/^{12}C$ HSI images of (A), (B) and (C), respectively. Note that some cell poles show Fe mineralization (blue arrows in C), and these regions show no $^{13}C$ accumulation (yellow arrows in F). The white arrow in (B) is indicating a possibly fully mineralized cell with relatively low levels of $^{75}As$.

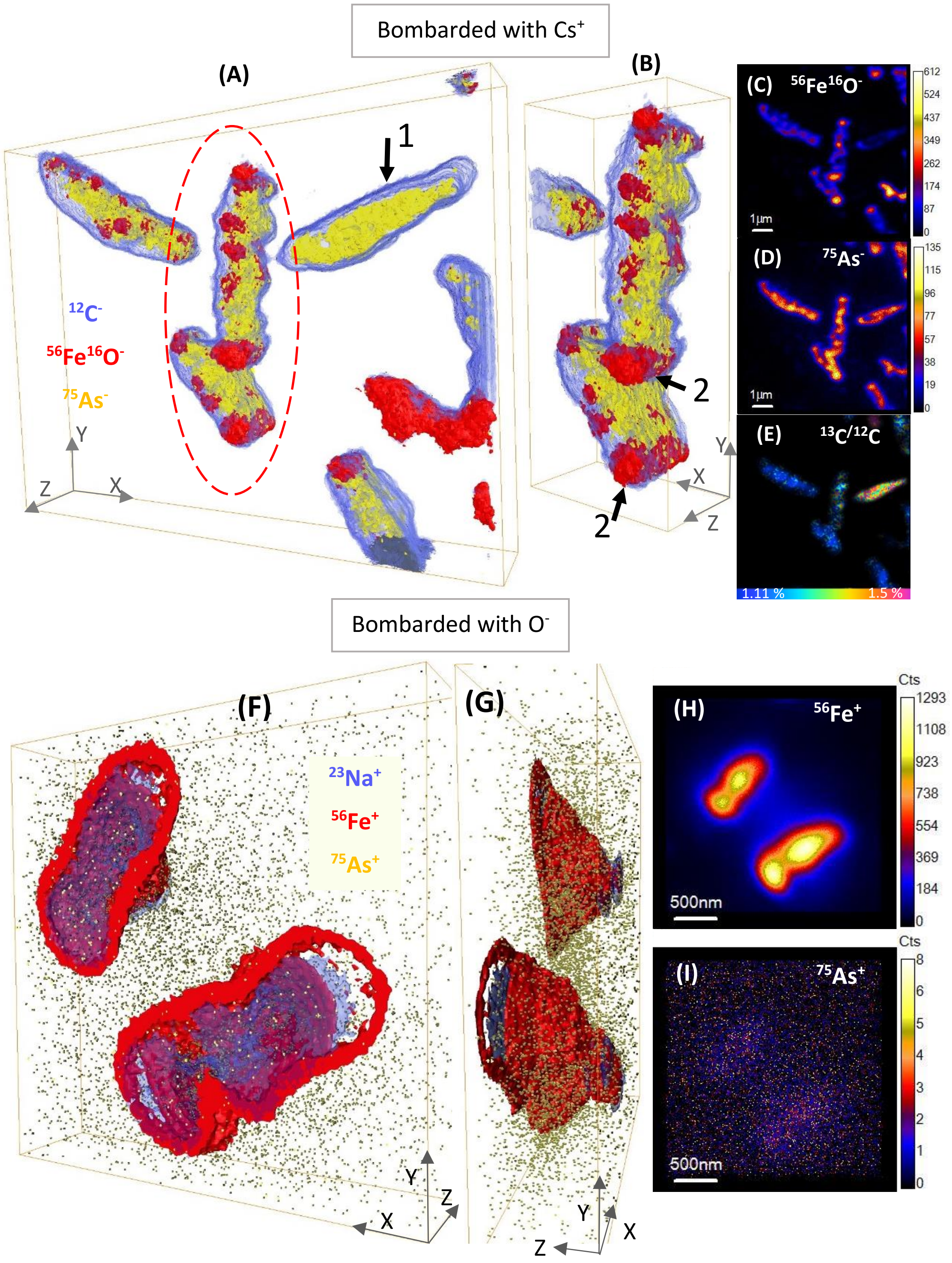


**Figure S6.** NanoSIMS depth profile 3D reconstructions of strain ST3 cells at day 7 of incubation. (A), (B), (F) and (G) are 3D reconstructions, where (A) & (B) were sputtered with $Cs^+$ (negative secondary ion mode), (A) is the whole area analyzed and (B) are the cells selected in the red dashed oval in (A) seen at another angle; (F) and (G) are cells sputtered with $O^-$ (positive secondary ion mode), where (G) is a side view of cells from panel (F). In these 3D reconstructions (A), (B), (F) & (G) Fe is shown in red (either $^{56}Fe^{16}O^-$ or $^{56}Fe^+$), $^{75}As$ in yellow and $^{12}C^-$ [(A) & (B)] or $^{23}Na^+$ [(F) & (G)] in blue. (C) and (D) are stack images of $^{56}Fe^{16}O$ and $^{75}As$, respectively, of the same cells in panel (A); (E) is the $^{13}C/^{12}C$ ratio in this area, sum of 170 planes. (H) and (I) are stack images of $^{56}Fe$ and $^{75}As$, respectively, of the cells in panel (F), sum of 101 planes. The arrow #1 in panel (A) is indicating a cell with no Fe encrustation but As on the surface, whereas arrows #2 in panel (B) are indicating the encrusted cell poles.

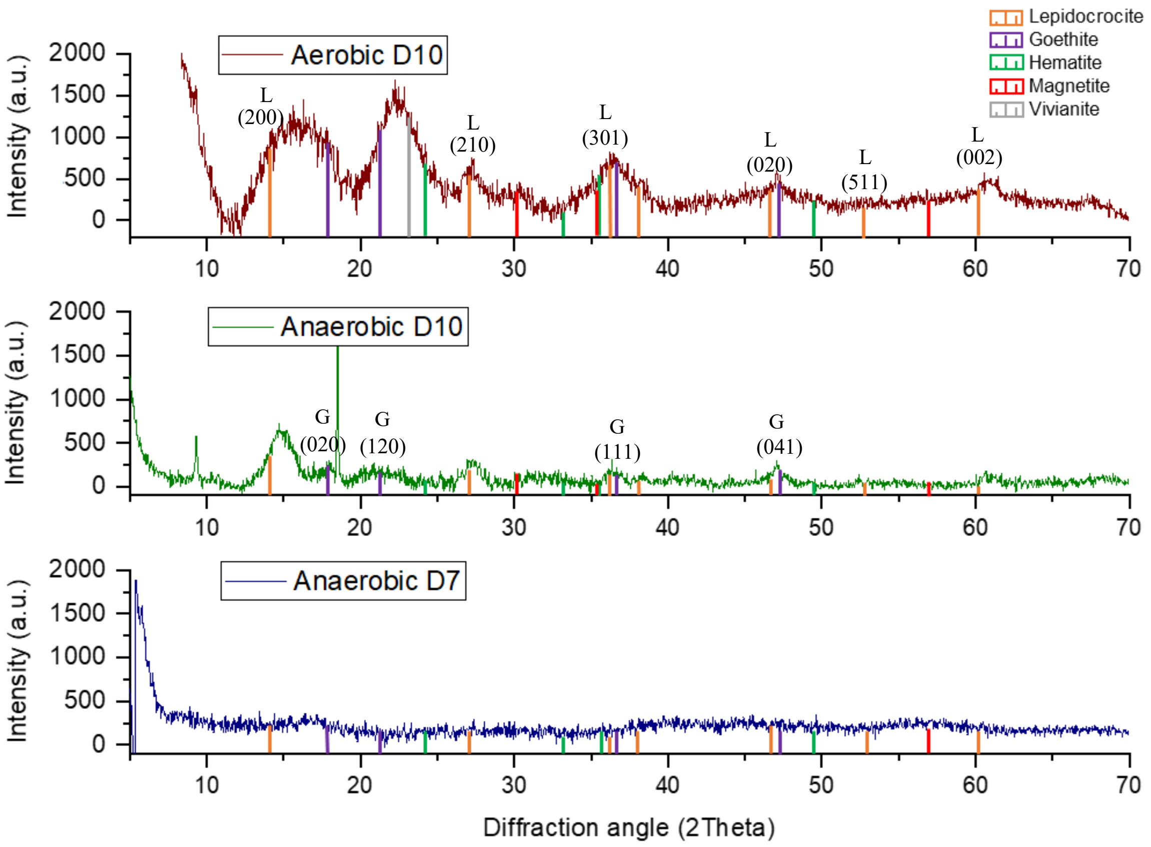


**Figure S7.** Diffractograms of the bulk precipitates of *Acidovorax* sp. strain ST3 at days 7 (anaerobic XRD analysis) and 10 of incubation (aerobic and anaerobic analysis). Aerobic analysis was collected without the air-tight dome, which produced higher intensity peaks. At day 10 lepidocrocite (L) and goethite (G) were the main mineral phases detected, more clearly noticed in the oxygen-exposed analysis, although peak matching indicates the presence of hematite, magnetite and vivianite.

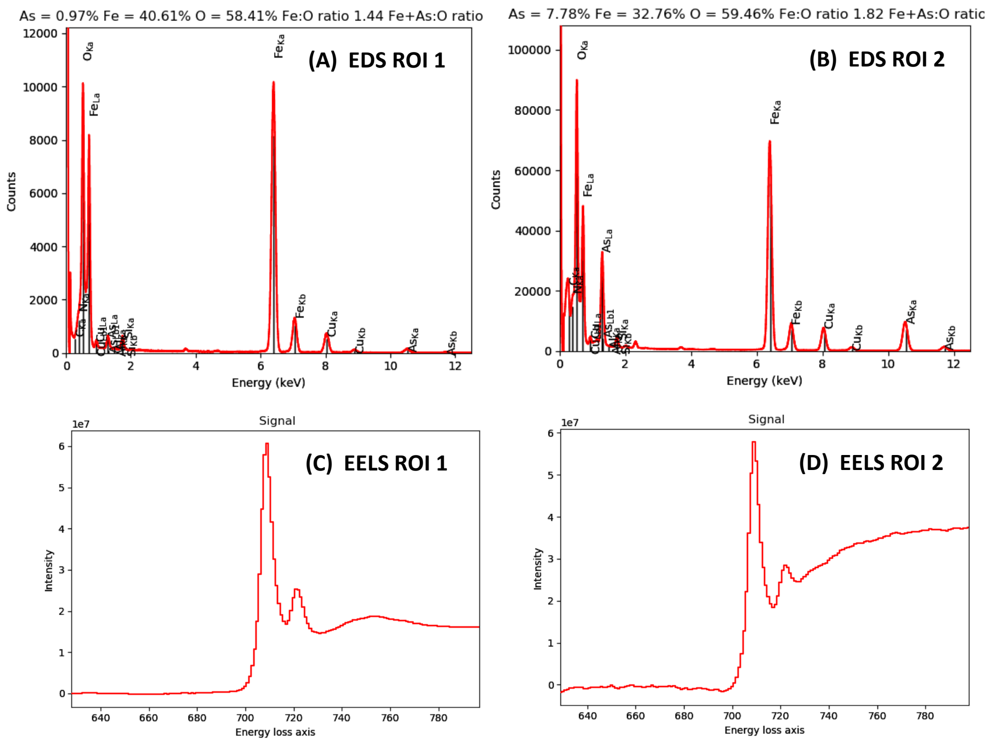


**Figure S8.** EDS and EELS spectra from the STEM high-angle annular dark field (HAADF) micrographs of two ROIs of mineralized *Acidovorax* sp. ST3 cells shown in Fig. 5. (A) and (B) are the EDS spectra of ROI 1 and 2, respectively, and (C) and (D) are EELS spectra of the same ROIs. EDS spectra show O and Fe are the most abundant elements in both ROIs, whereas the As peak varied in intensity; this intensity was higher in ROI 2 as well as its abundance. The $L_3/L_2$ intensity ratio was used to estimate $Fe^{3+}$ abundance and the EELS spectra show the $L_3$ and $L_2$ peaks.

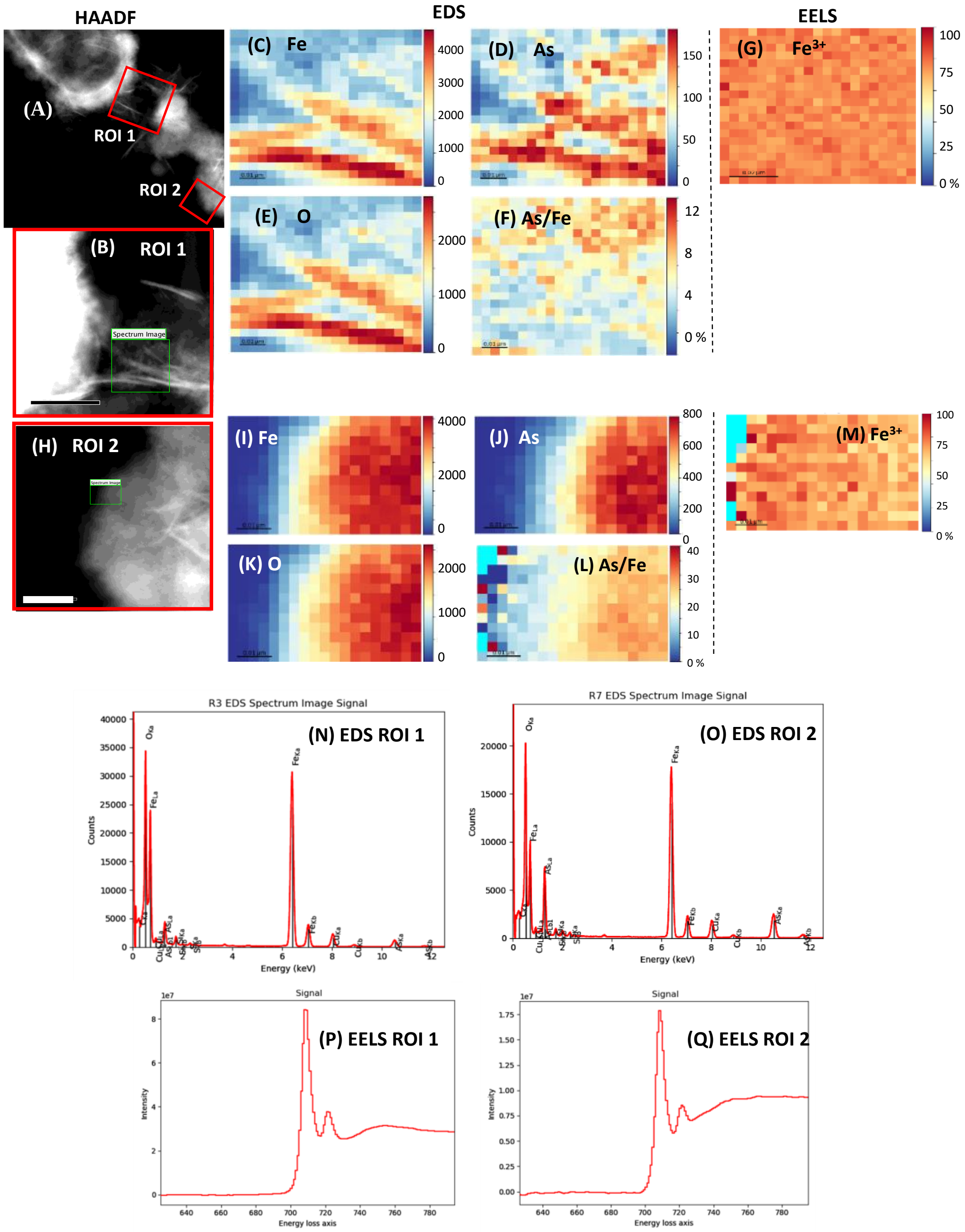


**Figure S9.** Additional STEM high-annular dark field (HAADF) micrographs with complementary EDS/EELS analysis of mineralized ST3 cells. Panel (A) is a HAADF micrograph of an area showing a region of the cell and extracellular biominerals, where regions of interest (ROI) were selected and further analyzed (red squares ROI 1 & 2); these ROIs are magnified in (B) & (H). (B-D) and (G-I) are EDS maps of the selected regions (green squares) in (B) and (H), respectively. (C) & (I) are maps of $^{56}Fe$, (D) & (J) are $^{75}As$ and (E) & (K) are $^{16}O$ maps. (F) and (L) are the As/Fe ratio maps (colour scale on the right, atomic %), where the average As/Fe ratio was 23.4 % in (F) and 6.7 % in (L). (G) & (M) are EELS maps of the $Fe^{3+}$ /$Fe^{2+}$ percentage composition, where $Fe^{3+}$ predominated, with an $Fe^{3+}$ average of 76 % in (G) and 72 % $Fe^{3+}$ in (M). Notice that Fe and O co-locate in both mapped regions in EDS. The scale bars are 100 nm for the images in (B) and (H). EDS spectra (N & O) show O and Fe are the most abundant elements in both ROIs, whereas the As peak varied in intensity; this intensity was higher in ROI 1 and so was its abundance. The $L_3/L_2$ intensity ratio was used to estimate $Fe^{3+}$ abundance and the EELS spectra (P & Q) show the $L_3$ and $L_2$ peaks.

### References


1. Newsome, L.; Lopez Adams, R.; Downie, H. F.; Moore, K. L.; Lloyd, J. R., NanoSIMS imaging of extracellular electron transport processes during microbial iron(III) reduction. *FEMS Microbiol Ecol* **2018,** *94* (8).
2. Wilson, R. G., SIMS quantification in Si, GaAs, and diamond-an update. *International Journal of Mass Spectrometry and Ion Processes* **1995,** *143* (C), 43-49.
3. Robinson, M. A.; Castner, D. G., Characterization of sample preparation methods of NIH-3T3 fibroblasts for ToF-SIMS. *Biointerphases* **2013,** *8* (15), 1-12.
4. Fearn, S., Characterisation of biological material with ToF-SIMS: a review. *Materials Science and Technology* **2015,** *31* (2), 148-161.
